# KIF5B and Dynein regulate adhesion-dependent Golgi organization and microtubule acetylation

**DOI:** 10.1101/2025.05.11.653231

**Authors:** Antara Chakraborty, Sharvari M. Pitke, BR Rajeshwari, Anwesha Dasgupta, Natasha Buwa, Rajalaxmi Behera, Madhavan Jayakrishnan, Nagaraj Balasubramanian

## Abstract

Cell-matrix adhesion regulates Golgi organization through Arf1-mediated dynein recruitment, maintaining its juxtanuclear localization. On loss of adhesion, Arf1 activation drops, causing loss of dynein, promoting differential disorganization of cis- vs trans-Golgi along microtubules. Golgi regulates microtubule nucleation and stability. In fibroblasts, acetylated tubulin levels drop on loss of adhesion, recovering on re-adhesion with time. Active Arf1 overexpression in preventing Golgi disorganization sustains microtubule acetylation, also seen in T24 bladder cancer cells. Active Arf1 binds KIF5B, recruiting it to the Golgi. KIF5B and dynein knockdown disorganize the Golgi as ministacks, with cis- and trans-Golgi. Dynein knockdown disrupts MTOC positioning, causing ministacks to disperse, preventing Golgi reorganization upon re-adhesion. Dispersed ministacks interestingly maintain microtubule acetylation in adherent and non-adherent cells. The joint KIF5B-dynein knockdown causes the Golgi to lose its ribbon morphology, becoming compact while keeping cis- and trans-Golgi together. This also causes a change in spreading, aspect ratio and migration of knockdown cells, which could be regulated by their Golgi phenotype. In evaluating adhesion-dependent Golgi organization, we reveal the Arf1-KIF5B-dynein crosstalk to regulate Golgi-dependent tubulin acetylation and cell function.

**Summary:** KIF5B and dynein are vital microtubule-associated motors that drive organelle positioning and organization. Adhesion-dependent Arf1 activation mediates KIF5B and dynein’s recruitment to the Golgi, regulating its organization and position. This, in turn, regulates microtubule acetylation levels, localization, and cellular functions.

## INTRODUCTION

Cells receive various mechanical and chemical cues from their extracellular environment. Cell-matrix adhesion is one of the most vital signalling cues cells respond to. Cell-matrix adhesion regulates signalling (Palazzo et al., 2004; Buwa et al., 2021), which, combined with its regulation of the cytoskeleton (Bershadsky et al., 1996; Small and Kaverina, 2003), controls cell division, survival, and function (Kaverina and Straube, 2011; Deakin and Turner, 2014; Guo et al., 2017). Cell organelles constitute important regulatory players in cellular functions, known to be regulated by adhesion (B.R. et al., 2023; Crosas-Molist et al., 2023; Singh et al., 2018). Cell-matrix adhesion-dependent regulation of Golgi organization is mediated by its regulation of the small GTPase Arf1 (Singh et al., 2018), and supported by the microtubule network. Active Arf1 and Golgin 160 recruit dynein to the Golgi, causing minus-end-directed movement of the Golgi, which helps keep its juxtanuclear localization (Yadav et al., 2012; Singh et al., 2018). Upon loss of adhesion, active Arf1 is differentially lost from the cis and the trans-Golgi, causing them to disorganize to varying extents. The trans-Golgi disorganizes more than the cis-Golgi (B.R. et al., 2023) which could have implications for the Golgi and cell function (B.R. et al., 2023). The different cisternae of the Golgi house glycosyltransferases and glycosidases, which, being in the correct order, are vital for sequential lipid and protein processing as they pass through the cisternae (Xiang et al., 2013; Pinho and Reis, 2015; Joshi et al., 2014, 2015). The Golgi complex regulates protein and lipid glycosylation, sorting, and trafficking. Golgi structural defects are observed in cells of several diseases, like Alzheimer’s disease (Huse et al., 2002; Joshi et al., 2014), Huntington’s disease (Hilditch-Maguire, 2000), and amyotrophic lateral sclerosis (Fujita and Okamoto, 2005). Cancer (Kellokumpu et al., 2002; Manca et al., 2019; Zhang, 2021), diabetes (Limoge et al., 2015), and COPD (Pavić et al., 2018; Weidner et al., 2018) also show Golgi organization and Glycosylation defects, which could be influenced by cell-matrix adhesion.

Microtubule acetylation is a post-translational modification (PTM) often found in stable, long-lived microtubules in the K40 position of alpha-tubulin (Reed et al., 2006; Janke and Montagnac, 2017). Cell-matrix adhesion is reported to regulate the levels of tubulin acetylation (Wen et al., 2023). Cells plated on lower matrix stiffness or non-adherent conditions have a lesser acetylated tubulin population than cells on higher stiffness (Ko et al., 2021; Wen et al., 2023). Focal adhesion scaffold protein paxillin inhibits HDAC6 activity, which regulates tubulin acetylation levels (Deakin and Turner, 2014). RNAi-mediated knockdown (KD) of paxillin causes a drop in microtubule acetylation and results in Golgi disorganization. A tubacin-mediated rise in microtubule acetylation rescues the Golgi phenotype in paxillin KD cells (Deakin and Turner, 2014). In turn, microtubule acetylation is also reported to control the expression of focal adhesion proteins and the dynamics of nascent focal adhesions (Ko et al., 2021).

Along with the levels of acetylated tubulin, the directionality and organization also play a vital role in regulating directional trafficking, polarity, and cell migration (Efimov et al., 2007). The localization and positioning of the Golgi are critical for controlling the directionality of acetylated tubulin in the cell (Rong et al., 2021). Golgi is an important microtubule-nucleating and organizing centre alongside the centrosomes (Sanders and Kaverina, 2015). Studies have also shown that centrosomes are dispensable for Golgi-derived nucleation of microtubules (Efimov et al., 2007). Cis-Golgi protein GM130 recruits AKAP450, a centrosomal γ-TuRC-interacting protein responsible for Golgi-derived microtubule nucleation (Rivero et al., 2009). Microtubule plus tip protein CLASP1 and CLASP2 (together called CLASPs), which localize to the trans-Golgi network, are also shown to be indispensable for Golgi-derived microtubule nucleation (Efimov et al., 2007). Trans-Golgi network protein GCC185 recruits CLASPs to the trans-Golgi network, and its depletion impairs microtubule nucleation at the Golgi and alters the microtubule pattern (Miller et al., 2009). CLASPs preferentially coat the plus-end tips of freshly nucleated Golgi-derived microtubules. CLASP coating stabilizes microtubule seeds on the Golgi, preventing them from depolymerization and promoting the growth of tubules from the Golgi (Efimov et al., 2007). The presence of AKAP450 and CLASP in the Golgi ministacks allows for microtubule nucleation and growth upon nocodazole washout, suggesting that cis and trans-Golgi proteins are sufficient for this process (Rivero et al., 2009; Efimov et al., 2007). This also suggests that the Golgi ribbon organization could be dispensable for Golgi-derived microtubule nucleation and polymerization.

Golgi ministacks are considered functional Golgi subunits (Okumura et al., 2023), where cis- and trans-Golgi compartments sit next to each other, supporting tubulin acetylation levels (Okumura et al., 2023). Loss of adhesion causes differential disorganization of the cis- and trans-Golgi (Singh et al., 2018; B.R. et al., 2023), which could regulate microtubule acetylation. The known role of dynein in regulating cell-matrix adhesion-dependent Golgi organization raises the possibility of a plus-ended kinesin acting alongside it. Arf1 has earlier been shown to recruit KIF5B to lipid droplets (Kumar et al., 2019; Rai et al., 2017). KIF5B localizes to the Golgi in cortical neuronal (Wobst et al., 2015) and HeLa cells (Mahajan et al., 2019). This led to our hypothesis that Arf1 could work to recruit dynein and KIF5B at the Golgi to regulate their cell-matrix adhesion-dependent organization along microtubules. Golgi-organization-dependent regulation of microtubules could further add to this crosstalk and have implications for how this affects cell function in normal and anchorage-independent cancer cells.

## MATERIALS

### Reagents

Fibronectin (Cat. No. # F2006), Accutase (Cat. No. # A6964), DMSO (Cat. No. # D2438), Triton-x 100 (Cat. No. #T8787), Nocodazole (Cat. No. #M1404), and Tubacin (Cat. No. #SML0065) were purchased from Sigma. Ciliobrevin D (Cat. No. # 250401) was purchased from Calbiochem.

Fluoromount-G (Cat. No. #0100-01) was purchased from Southern Biotech. Immobilon Western Chemiluminescence substrate was purchased from Millipore (Cat. No. #WBKLS0500). BCA protein estimation kit (Cat. No. #23225) was purchased from Pierce. DAPI was purchased from Merck (Cat. No. #5.08741.0001). Phalloidin was purchased from Invitrogen (Cat. No. #A12379, A12381, A22287). Fluorophore-conjugated lectin probes were purchased from Molecular Probes - ConA-Alexa647 (Cat. No. #C21421), WGA-Alexa488 (Cat. No. #W11261).

### Plasmids and oligos

GFP-tagged Arf1-WT construct was obtained from Dr. Satyajit Mayor (National Centre for Biological Sciences, Bangalore, India). GFP-tagged Arf1-Q71L construct was made as earlier described (Singh et al., 2018). Galtase-RFP (GalT-RFP) was obtained from Dr. Richa Rikhy (IISER Pune, originally from Dr. Jennifer Lippincott, NIH). Mannosidase II-GFP (ManII-GFP) was obtained from Dr. Jennifer Lippincott (NIH). mCherry-tagged Arf1-WT, T31N-Arf1, Arf1-Q71L constructs were made as earlier described (Singh et al., 2018). All of the constructs mentioned above were used after the sequence was confirmed.

siRNAs were purchased from Sigma oligos. The following siRNA duplexes were used: scramble scrKIF5B: 5’-AGACGACUAACUCAGAUUG-3’, scramble scrDynein Heavy Chain: 5’-AUUCCUUCAAUACCAUACA-3’, siKIF5B: 5’-GACAUGUCGCAGUUACAAA-3’, siDHC (Dynein Heavy chain): 5’-CCAAAUACCUACAUUACUU-3’. All sequences were checked and confirmed for target specificity using the NCBI nucleotide BLAST tool.

### Antibodies

| Antibody | Catalog number | Dilution used |
| --- | --- | --- |
| Anti-GM130 | BD transduction, Clone 35, Cat. No. #610822 | 1:500 for WB and IFA |
| Anti-Giantin | Abcam, Cat. No. #ab24586 | 1:200 for IFA |
| Anti-Dynein | Millipore, Clone 74.1, Cat. No. #MAB1618 | 1:2000 for WB |
| Anti-KIF5B | Abcam, Cat. No. #ab5629 | 1:2000 for WB,<br>1:200 for IFA |
| Anti-Calnexin | Abcam, Cat. No. #ab10286 | 1:2000 for WB |
| Anti-Rho-GDI | Millipore, Cat. No. #6730 | 1:2000 for WB |
| Anti-GAPDH | Sigma, Cat. No. #G9545 | 1:5000 for WB |
| Anti-Beta-Actin | Abcam, Clone ACTN05(C4), Cat. No. #ab3280 | 1:2000 for WB |
| Anti-Arf1 | Abcam, Cat. No. #ab183576 | 1:3000 for WB |
| Anti-Beta tubulin | Developmental Studies Hybridoma Bank, Clone E7, Cat. No.#AB_2315513 | 1:2000 for WB<br>1: 1000 for IFA |
| Anti-Alpha tubulin | Developmental Studies Hybridoma Bank, Clone 12G10, Cat. No. #AB_1157911 | 1: 2000 for WB<br>1:1000 for IFA |
| Anti-Acetylated tubulin | Sigma, Clone 6-11B-1, Cat. No. #T7451 | 1: 2000 for WB<br>1:1000 for IFA |
| Anti-Detyrosinated tubulin | Adipogen Life Sciences, Clone RM444, Cat. No. #REV-31-1335-00-R100 (Kind gift from Prof. Aurnab Ghose, IISER Pune) | 1: 5000 for WB |
| Anti-γ-tubulin | Abcam, Cat. No. #ab11316 | 1:100 for IFA |
| HRP-tagged Secondary antibodies | Jackson Immuno Research | 1:5000 for WB |
| Fluorophore-tagged Secondary antibodies | Invitrogen Molecular Probe | 1:1000 for IFA |

## METHODS

### Cell culture and transfections

Wild type- mouse embryonic fibroblasts (WT-MEFs), obtained from Dr. Richard Anderson (University of Texas Health Sciences Center, Dallas, TX), and T24 bladder cancer cells, obtained from ECACC, were cultured in complete DMEM medium (Invitrogen) supplemented with 5% fetal bovine serum (FBS) (Invitrogen). Cells were grown at 37 °C in a 5% CO_2_ humidified atmosphere. According to the manufacturer’s protocol, cells were transfected with different DNA constructs using Lipofectamine 2000 (Invitrogen). Transfections were done in 6 cm plates using 4 µg DNA per well, and the transfection mixture was removed after 12 hours. Cells were used for suspension experiments 48 hours post-transfection. For all the experiments, MEFs were serum-starved for 14 hours before suspension assay in DMEM medium with 0.2% FBS.

### siRNA-mediated KIF5B KD, Dynein KD, and Dual KD

siRNA-mediated knockdowns (KD) were done using Lipofectamine RNAiMax (Invitrogen). WT-MEFs (0.6 x10^5^) seeded in 3.5 cm dishes for 3-4 hours treated with scramble or siRNA constructs against Kinesin-1 (KIF5B) (50 picomoles), Dynein Heavy Chain (100 picomoles), or both. RNAiMax with/without scrambled siRNA-treated cells were used as controls. This treatment was repeated 24 hours after the first shot. After 48 hours, cells were serum-deprived (0.2% FBS DMEM) for 14 hours, detached, counted, and used for various assays.

### Inhibitors studies

15 x 10^5^ WT-MEFs were seeded in 10 cm dishes for ∼24 hours; cells were serum-deprived with low-serum DMEM (0.2% FBS) with 1 µM Tubacin or DMSO for 14 hours or 100 µM ciliobrevin D or DMSO for 2 hours. For both treatments drug was added at the same concentrations in suspension and re-adhesion of cells. Control and dual KD cells were treated with 10 µM Nocodazole for 40 minutes or equivalent DMSO. Following drug treatment, cells were processed for Arf1 activity assay or immunofluorescence staining and imaging as described later.

### Suspension and re-adhesion assay

80% confluent cells were serum starved (0.2% FBS) for 14 hours for MEFs, or maintained in DMEM with 5% FBS for T24, detached with Trypsin-EDTA, and the required number of cells were suspended in 0.8% methylcellulose for 120 minutes. Post-incubation cells were handled carefully to avoid clumping. Methylcellulose was washed with media twice at 4 °C to harvest cells. Aliquots of cells were collected post-suspension, and for re-adhesion, cells were seeded on 22 x 22 mm coverslips or 6 cm dishes pre-coated with 10 μg/ml fibronectin. When treated with inhibitors, they were added to the suspension mix, followed by media washes, and re-adhesion timepoints. For confocal fluorescence microscopy, cells were fixed with 3.5% paraformaldehyde for 10 minutes at room temperature (RT) and/ or chilled methanol for 15 minutes at -20 °C, washed with PBS thrice, and processed. For the Arf1 activity assay, cells were lysed in an activity assay buffer followed by GST-GGA3 pulldown. For western blotting, cells were lysed in Laemmli buffer, and boiled at 95 °C for 15 minutes.

### Golgi-enriched membrane fraction isolation

Total membrane isolation was done from adherent cells. Adherent cells grown in 15 cm dishes were scraped in 5 ml of PBS and spun down at 200 g for 5 minutes. Cell pellets were washed with 5 ml PBS, then 1 ml of homogenising buffer to remove traces of PBS. All these steps were done on ice or at 4 °C. Cells were homogenized with 65-70 strokes in a Dounce homogeniser. Samples were then processed through sequential centrifugation at 4 °C. The sample was centrifuged at 200 g for 3 minutes to pellet cell debris. The supernatant was centrifuged at 1000 g for 15 minutes to pellet the nuclear fraction. The post-nuclear supernatant was collected and centrifuged at 10,000 g for 15 minutes to pellet the mitochondrial fraction. The supernatant was centrifuged at 100,000 g for 1 hour in a TLA 100.3 rotor in a tabletop ultracentrifuge to pellet the membrane fraction. The supernatant here is the cytoplasmic fraction. 30 µl of the cytosolic and membrane fractions were used for protein estimation, and the rest mixed with Laemmli buffer and boiled at 95 °C for 15 minutes. These fractions were analyzed by western blotting.

### Protein estimation with Bicinchoninic acid assay (BCA) kit

Samples lysed in RIPA buffer with protease inhibitors and kept on ice for 30 minutes were spun down at 14000 rpm; 4 °C for 15 minutes. The supernatant was collected in a fresh tube as the lysate. For protein estimation in a 96-well plate, samples were diluted 1:5 and 1:10 times with RIPA buffer (in triplicates) along with freshly prepared BSA standards diluted in RIPA buffer (0mg/ml to 2mg/ml). Working reagent from the Pierce BCA kit was prepared by mixing Reagent A and Reagent B in a ratio of 50:1. 200 µl of working reagent was added to each well, containing 10 µl of sample or BSA standard. The plate was incubated at 37° C for 30 minutes, and absorbance was determined at 562 nm using a plate reader (PerkinElmer Ensight). BSA absorbance values were plotted to obtain a standard curve, determining the unknown sample’s protein concentration.

### Arf1 activity assay

6 x 10^5^ cells per time point (suspension or adherent), were lysed in 500 μl Arf activity assay buffer (Pawar et al., 2016; Singh et al., 2018) . 400 μl of this lysate was incubated with 60 μg of glutathione S-transferase (GST)-tagged Golgi-localized γ-ear containing Arf-binding protein 3 (GGA3) fusion protein (GST-GGA3) for 35 minutes, as described earlier(Singh et al., 2018; Pawar et al., 2016). Active Arf1 pulled down with the beads was washed and eluted in 20 μl of Laemmli buffer (PD). 100 μl of the whole-cell lysate (WCL) was kept aside and added to the Laemmli buffer. PD and WCL fractions were boiled at 95 °C for 15 minutes, and western blotting detection of Arf1 was done. Active Arf1 in the PD was normalized to its respective levels in the WCL. Proteins bound to the active Arf1 PD were also detected by western blot.

### Tubacin treatment and Arf1 activity assay

The same protocol was followed as above, with some changes. 15 x 10^5^ WT-MEFs seeded in 10 cm dishes were serum-deprived with 0.2% FBS DMEM containing either 1µM Tubacin or volume equivalent of DMSO for 14 hours. Following this, cells were detached, and 6 x 10^5^ cells were used for each time point, control, and treated conditions. Following suspension assay cells were washed with cold low-serum DMEM with drug or DMSO, centrifuged at 1000 rpm for 5 minutes at 4 °C, and reconstituted in low-serum medium with drug or DMSO. Cells were replated on FN-coated dishes for 15 minutes and 4 hours, washed, frozen, and lysed in the activity assay buffer. Active Arf1 was pulled down with GST-GGA3 fusion protein and detected as described above.

### Detection of KIF5B in GGA3 pulldown

Active Arf1 was pulled down using GST-GGA3 beads as described above. PD and WCL resolved by SDS-PAGE for western blot detection of Arf1 and KIF5B. GST-GGA3 bound protein normalized to their respective levels in WCL.

### SDS-PAGE and Western Blotting

The required protein or cell equivalent lysate was loaded on SDS-PAGE gels. Protein samples run on gels were transferred to the PVDF membrane (Pawar et al., 2016; Singh et al., 2018; B.R. et al., 2023). PVDF membrane was blocked in 5% skimmed milk (made in TBS-Tween20 (0.1%)) at RT for 1 hour. The membrane was washed with TBST; individual blots were incubated in respective primary antibody diluted in 5% BSA in TBST overnight at 4 °C. Blots were washed thrice with TBST, incubated at RT with respective HRP-conjugated secondary antibody diluted in 2.5% skimmed milk in TBST. Washed thrice with TBST, blots were developed with Immobilon substrate (diluted when needed with TBST). Blots were imaged using the GE Healthcare LAS4000 Chemiluminescent Imager. Densitometric analysis was done using ImageJ (NIH) software.

### Immunofluorescence

Cells plated on coverslips coated with 10 μg/ml FN-coated glass (re-adherent time points) or in a reconstituted pellet (suspension time points) were fixed with 3.5% paraformaldehyde (PFA), for 10 minutes at RT (Golgi staining) or chilled methanol for 15 minutes at -20 °C (acetylated or alpha tubulin staining), or a combination of both (PFA followed by Methanol), for Golgi and tubulin dual staining. Adherent cells in coverslips or suspended cells in eppendorf tube were permeabilized with 0.5% Triton-X 100 for 10 minutes at RT. Blocked with 5% BSA for 1 hour at RT. Incubated with appropriately diluted primary antibody overnight, followed by secondary antibody for 1 hour at RT. When needed cells were stained with 1:500 diluted Phalloidin Alexa Fluor 488 overnight at 4 °C. Cells were washed with PBS (10,000 rpm/5 minutes/4 °C for suspension time points) after each step and mounted using Flouramount-G. All slides were allowed to dry for 24 hours and imaged with a confocal microscope.

### Cell surface lectin binding and quantitation by flow cytometry

WT-MEFs were serum deprived for 14 hours, detached using Accutase, washed with 1X PBS, and incubated with dual-labelled with ConA-Alexa647, 5 ng/µl; WGA-Alexa488, 0.5 ng/µl. The labelling reaction was incubated on ice in the dark for 15 minutes, followed by two cold PBS washes. For flow cytometry, the lectin-labelled cells were fixed with 3.5% PFA for 10 minutes at RT, and resuspended in 350 µl of cold PBS. Data acquisition was performed using a BD Celesta Flow Cytometer, with a morphologically consistent cell population selected based on forward and side scatter (FSC and SSC) profiles using polygon gating.

### Imaging, cell spread, and aspect ratio analysis

Stained with Phalloidin, re-adherent WT-MEFs were imaged using EVOS FL Auto Imaging System (ThermoFisher Scientific) using 20x objective and analyzed using ImageJ (NIH). Acquired images were converted to 8-bit. A threshold is set to cover the entire stained cell area, defining the cell boundary. The wand (tracing) tool was then used to select cell boundaries. The area within this boundary and the cell aspect ratio were calculated using the “measure” option under the “Analyze” tab in ImageJ (NIH). These were measured for 250 or more cells for each treatment and timepoint to calculate means, which were plotted with individual values and compared across treatments.

### Single-cell migration assay

WT-MEFs were seeded on either glass-bottom dishes, or Lab-Tek chambered coverglass (Thermo Fisher Scientific) coated with 2 μg/ml FN, and allowed to spread for 12 hours. Cells were then imaged on the heated stage for an additional 12 hours, with images taken at 10-minute intervals in DIC mode, on an Olympus SpinSR Spinning Disk Confocal Microscope with a 10x objective. Cells were maintained in 5% FBS DMEM, 37 °C in a 5% CO_2_ humidified atmosphere. For migration analysis, TrackMate (Ershov et al., 2022) was used in ImageJ (NIH) to map and track cells. This was used to determine total distance travelled, displacement, and confinement ration for individual cells tracked for the entire length of imaging.

### Confocal Microscopy

Images were acquired using Zeiss 710 and 780 laser scanning confocal microscopes with a 63× oil objective (NA 1.4). Acquisition settings were laser power=2%, Pinhole=1 AU, gain=700–950. Images were acquired at a resolution of 1024×1024, with a scan speed of 5 for cross-sections and 7 for Z-stacks. Images are shot at variable zoom; scale bars are denoted in every image. Z-stacks were acquired at 0.2 µm intervals, deconvoluted, and rendered when needed.

### Deconvolution of Z-stacks and object analysis

Images were processed and analyzed using the Huygens Professional version 16.10 (Scientific Volume Imaging, The Netherlands, http://svi.nl). Deconvolution of confocal Z-stacks was done at average background value =1, number of iterations =30, signal-to-noise ratio (SNR) =20, and quality change threshold =0.0001. These were kept constant for deconvolutions. Point spread function (PSF) values were estimated for each cross-section or Z-stack, provided the minimal voxel size the confocal microscope could resolve. This minimum voxel value was used to calculate garbage volume for object analysis. Deconvoluted images were surface-rendered. Surface-rendered images were pseudocoloured when needed. The number of discontinuous Golgi objects in a 3D deconvoluted image was determined using the advanced object analysis plugin. The garbage volume was set as calculated earlier, and an optimal threshold was used to determine the number of discontinuous Golgi objects present in a cell.

### Number of objects

Golgi object number was calculated as the number of discontinuous voxels present in a deconvoluted 3D image after setting an optimal threshold. An object is defined only when it has several voxels more than the point spread function of that image. Otherwise, it was considered noise and was subtracted as garbage volume. The number of objects for each cell was noted, and the average number of Golgi objects was compared between different conditions.

### Colocalization analysis

De-convoluted Z-stack images were opened with the Colocalization Analyzer plug-in in the SVI Huygens Professional software (version 16.10), and Pearson coefficients were calculated for each cell. Costes’ method was used to set a threshold for each image, and the Golgi channel was manually thresholded and analyzed by the software to obtain Pearson’s coefficient value.

### Distribution profile of Golgi organization and MTOC localization

Two Golgi morphologies (organized and disorganized) were used to define a distribution profile. Three MTOC positions (proximal, in-between, and away with respect to their distance from the nucleus) were used to define a distribution profile. A minimum of 200 cells was counted and represented as a percentage distribution profile. Cross-sectional images and/or Z-stacks for each morphology were acquired for each time point and treatment in every experiment. Percentage distribution profiles were compared, and differences were evaluated.

### Microtubule architecture analysis

Confocal Z-stack images for microtubules were deconvoluted using Huygens Professional software. Using Image J (NIH), batch processing was done using a custom macro written in the lab to apply standardised image analysis steps, ensuring consistency across all datasets. Before processing, cells were selected through manual thresholding by marking an ROI, and the background signal was eliminated by running the command ‘Clear Background’. The analysis protocol is adapted from an earlier study focusing on mitochondrial architecture (Chaudhry et al., 2020). The sequential steps in the macro include (1) background subtraction (rolling = 15.17), (2) sigma filter plus (radius = 1.138), (3) CLAHE (block size = 64, max slope = 1.25), (4) gamma correction (value = 0.90), (5) adaptive thresholding (Mean from = 15, then = -27). The function ‘Despeckle’ and ‘outlier removal’ (block radius x/y = 1.707, standard deviation = 3) were also applied to reduce noise and enhance image features. Processed Z-stacks were skeletonized using the *Skeletonize (3D)* plugin to create a binary skeleton image. Skeletonized stacks were analyzed using the *Analyze Skeleton* 2D/3D plugin (Lee et al., 1994). The Analyze Skeleton plugin measured the total number of branches and average branch length in the skeletonized image (Arganda-Carreras et al., 2010). The total branch length was calculated by summing the product of the average branch length and the branch number.

### Statistical analysis

All the analyses were done using Prism GraphPad analysis software. The Mann-Whitney t-test was used to statistically analyse the number of Golgi objects, Arf1 activation assay (GGA3 pulldown), Pearson coefficient, and western blot (non-normalized). When the data were normalized to a control and compared, a one-sample t-test was used. The one-way ANOVA test was used to statistically analyse changes in the distribution profile of the Golgi phenotype.

## RESULTS

### Cell-matrix adhesion-dependent regulation of stable microtubule levels: Cause or effect of adhesion-dependent Golgi organization?

Cell-matrix adhesion regulates Golgi organization (Singh et al., 2018). Golgi organization is vital for regulating microtubule stability, which can be determined by its PTMs like acetylation of lysine 40 on alpha-tubulin and proteolytic removal of tyrosine from the C-terminal of alpha-tubulin (Wloga and Gaertig, 2010). Adhesion-dependent cellular functions are regulated by microtubules (Deakin and Turner, 2014; Small and Kaverina, 2003; Bershadsky et al., 1996). A cause-and-effect relationship between adhesion-dependent Golgi organization and microtubule stability could exist. To test this, stable adherent wild-type mouse embryonic fibroblasts (WT-MEFs) grown in low-serum conditions for 14 hours to suppress growth factor signalling, were detached, held in suspension (with methylcellulose), and re-plated on fibronectin (FN) for early (15 minutes) and later (4 hours, stable adherent) time points. Upon loss of adhesion, microtubule acetylation levels drop by ∼50%, which does not recover on early re-adhesion on western blot (**Fig. 1 A)** and immunofluorescence detection **(Fig. 1 B)**. Time kinetics evaluation of how long it takes suspended cells to recover their microtubule acetylation and detyrosination levels on re-adhesion revealed a gradual increase over 4 hours **(Fig. 1 C)**, which could further increase over time.

**Figure 1:**
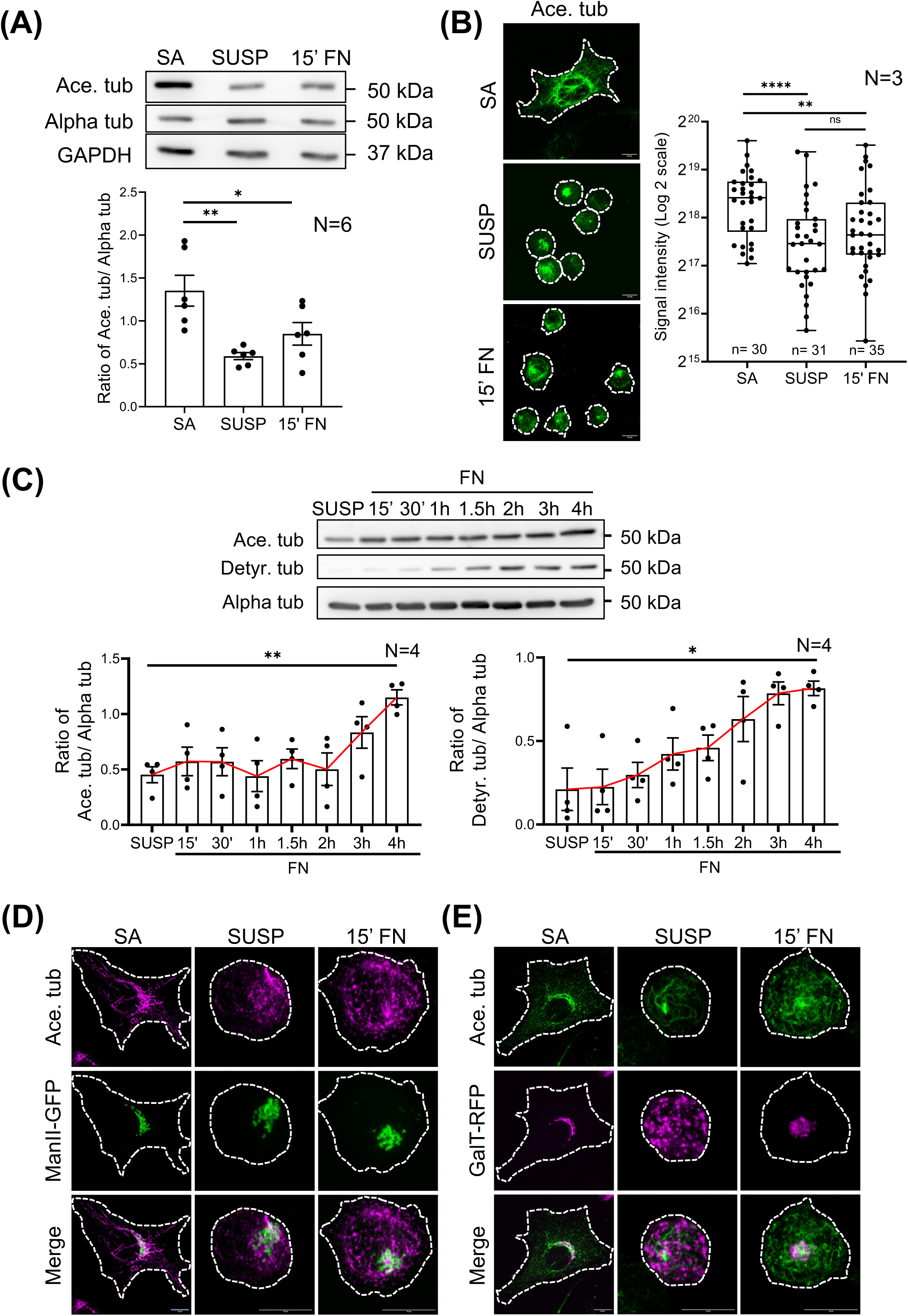
Cell-matrix adhesion-dependent regulation of Microtubule stability: **(A)** Western blot detection of Acetylated tubulin (Ace. tub), Alpha-tubulin (Alpha tub.), and GAPDH in the whole-cell lysate from stable adherent (SA), suspended (SUSP), and re-adherent (15′ FN) WT-MEFs. The graph represents ratio of densitometric band intensities as mean±SE from 6 independent experiments. **(B)** Collapsed Z-stack images from stable adherent (SA), suspended (SUSP), and re-adherent (15′ FN) WT-MEFs immuno-stained for Acetylated tubulin (Ace. tub). The graph represents fluorescence signal intensity as median and quarters from 3 independent experiments. **(C)** Western blot detection of Acetylated (Ace. Tub), Detyrosinated tubulin (Detyr. Tub), and Alpha tubulin (Alpha tub) in whole-cell lysate from suspended (SUSP) and re-adherent cells for increasing time points (15′, 30′, 1h, 1.5h, 2h, 3h, 4h). The graphs represent ratio of densitometric band intensities as mean±SE from 4 independent experiments. The red line connecting the means captures the trend observed **(D, E)** Collapsed Z-stack images from stable adherent (SA), suspended (SUSP), and re-adherent (15′ FN) WT-MEFs expressing **(D)** ManII-GFP or **(E)** GalTase-RFP, immunostained for Acetylated tubulin (Ace. tub). Scale bars-10 µm. Statistical analyses were done using Mann-Whitney’s t-test (non-normalized graphs) (* p-value<0.05, ** p-value <0.01, *** p-value <0.001, **** p-value <0.0001, ns = not significant).

Immunostaining for acetylated tubulin reveals a pronounced enrichment only in the juxtanuclear region adjacent to the cis-**(Fig. 1 D)** and trans-Golgi **(Fig. 1 E)**. On loss of adhesion, acetylated tubulin staining is retained near the cis-Golgi, which disorganizes less **(Fig. 1 D)** than the trans-Golgi, which is significantly more disorganized **(Fig. 1 E)**. On early re-adhesion, both the cis- and trans-Golgi reorganize rapidly, causing some accumulation of acetylated tubulin near the re-organized Golgi **(Fig. 1 D, E)**. This, we think, improves with time as acetylated tubulin levels also increase **(Fig. 1 C)**.

Earlier studies have shown that the expression of the focal adhesion scaffold protein, Paxillin inhibits HDAC6 to regulate microtubule acetylation (Deakin and Turner, 2014). RNAi-mediated depletion of paxillin causes a drop in tubulin acetylation, leading to Golgi disorganization in MDA-MB-231, Hs578T, and HFF cells (Deakin and Turner, 2014). We asked if changing paxillin levels could contribute to the loss of adhesion-mediated drop in microtubule acetylation and resulting Golgi disorganization. Western blot detection showed no significant change in paxillin levels on loss of adhesion and early re-adhesion (Supp Fig. 1 A), suggesting this regulation of the Golgi and microtubule acetylation could be independent of paxillin levels.

Golgi is known to nucleate microtubules with the help of AKAP1 and CLASP1/2, located on the cis and trans-Golgi network, respectively (Rivero et al., 2009; Efimov et al., 2007; Miller et al., 2009). Brefeldin A-mediated disruption of the Golgi network disrupts the organization of the microtubule network (Rivero et al., 2009). Arf1 KD leads to an increase in microtubule acetylation, as reported in oocytes during meiosis (Zhang et al., 2024). Microtubule acetylation levels are also shown to regulate Golgi organization (Ryan et al., 2012; Deakin and Turner, 2014). This suggests that loss of adhesion-mediated Golgi disorganization and a drop in microtubule acetylation could have a cause-and-effect relationship. To test this, we first prevented the drop in microtubule acetylation upon loss of adhesion to test if that affects Golgi disorganization. Second, does keeping the Golgi organized upon loss of adhesion prevent the drop in microtubule acetylation?

WT-MEFs were first treated with HDAC6 inhibitor Tubacin, raising microtubule acetylation levels and preventing a drop on loss of adhesion **(Fig. 2 A)**. The cis (Upper Panel) and trans-Golgi (Lower Panel) were disorganized in Tubacin-treated suspended cells, like DMSO-treated controls **(Fig. 2 B)**. Further, the loss of adhesion-mediated drop in active Arf1 levels is also unaffected by tubacin **(Fig. 2 C)**. This suggests that Arf1 activation and Golgi organization, regulated by cell-matrix adhesion, could act upstream of microtubule acetylation. To test this, constitutively active Arf1 (Q71L) was expressed in cells to prevent Golgi disorganization upon loss of adhesion (B.R. et al., 2023). Q71L expression prevented the drop in microtubule acetylation upon loss of adhesion **(Fig. 2 D, E)**. The basal microtubule acetylation is higher in cells expressing Q71L-Arf1 than WT-Arf1 **(Fig. 2 E)**. This suggests that Arf1 activation could promote microtubule acetylation levels independent of adhesion and Golgi organization. This is further substantiated in T24 cells with a compact, organized Golgi in stable adherent cells that remain organized upon adhesion loss. In T24 cells, the cis and trans-Golgi sit together when cells are adherent or suspended **(Fig. 2 F)**. Their active Arf1 levels do not drop on loss of adhesion **(Fig. 2 G),** which could prevent the Golgi from disorganizing upon loss of adhesion. Microtubule acetylation also does not drop on loss of cell-matrix adhesion **(Fig. 2 H)**. This reinforces our hypothesis that active Arf1 and Golgi organization could regulate microtubule acetylation levels downstream of cell-matrix adhesion.

**Figure 2:**
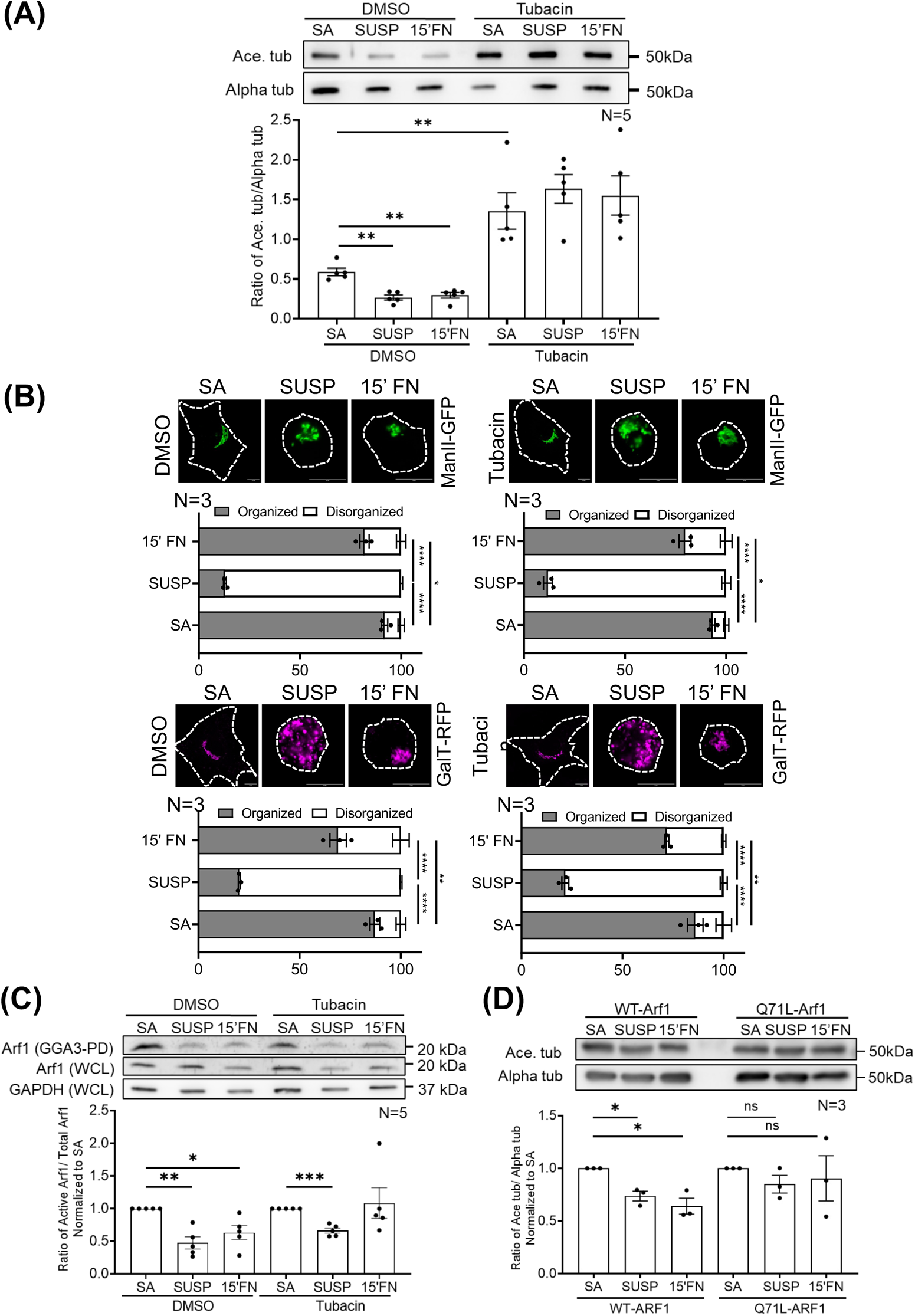
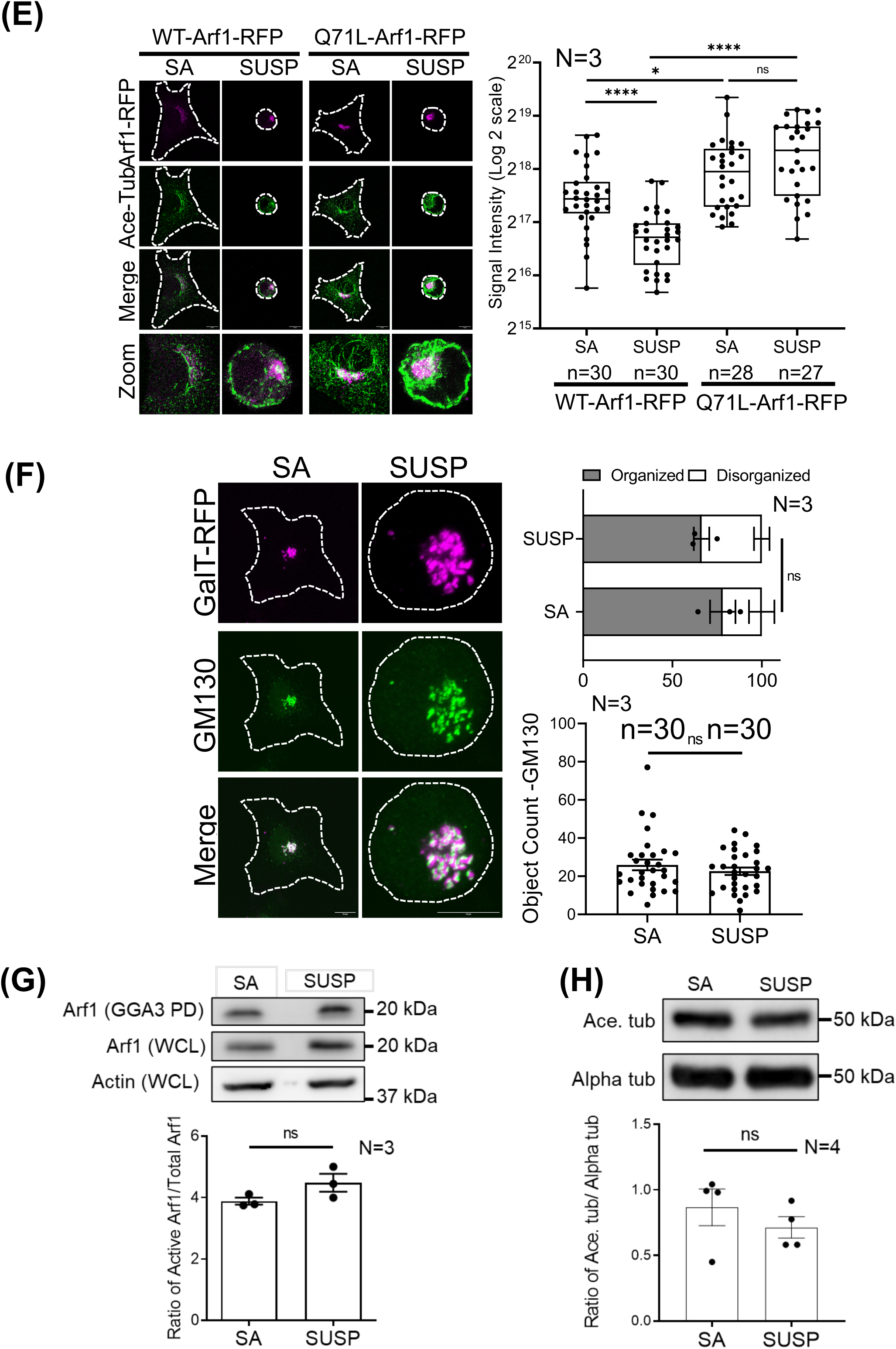
Cell-matrix adhesion-dependent drop in MT-Acetylation: Cause or effect of Golgi disorganization: **(A)** Western blot detection of Acetylated tubulin (Ace. tub) and Alpha tubulin (Alpha tub.) from DMSO or Tubacin-treated whole-cell lysate from stable adherent (SA), suspended (SUSP), and re-adherent (15′ FN) WT-MEFs. The graph represents ratio of densitometric band intensities as mean±SE from 5 independent experiments. **(B)** Cross-section images of DMSO or Tubacin-treated stable adherent (SA), suspended (SUSP), and re-adherent (15′ FN) WT-MEFs expressing ManII-GFP or GalT-RFP. The percentage distribution of cells with organized and disorganized Golgi phenotypes in these populations was determined and represented in the graph as mean±SE from 3 independent experiments. **(C)** Western blot detection of Arf1 in GST-GGA3 pulldown (GGA3 PD), whole-cell lysate (WCL), and GAPDH (WCL) from stable adherent (SA), suspended (SUSP), and re-adherent (15′ FN) WT-MEFs. The graph represents ratio of densitometric band intensities as mean±SE from 5 independent experiments. **(D)** Western blot detection of Acetylated tubulin (Ace. tub) and Alpha tubulin (Alpha tub.) in whole-cell lysate from stable adherent (SA), suspended (SUSP), and re-adherent (15′ FN) WT-MEFs expressing WT-Arf1 or Q71L-Arf1. The graph represents ratio of densitometric band intensities as mean±SE from 3 independent experiments. **(E)** Collapsed Z-stack images of stable adherent (SA) and suspended (SUSP) WT-MEFs expressing WT-Arf1-RFP or Q71L-Arf1-RFP, immuno-stained for Acetylated tubulin (Ace. tub). The graph represents fluorescence signal intensity as median and quarters from 3 independent experiments. **(F)** Collapsed Z-stack images of stable adherent (SA) and suspended (SUSP) T24 cells expressing GalT-RFP, immuno-stained for GM130. The above graph shows the percentage distribution of cells with organized and disorganized Golgi phenotypes in these populations. The graph below represents discontinuous Golgi object count per cell. Both graphs are represented as mean±SE from 3 independent experiments. **(G)** Western blot detection of Arf1 in GST-GGA3 pulldown (GGA3 PD) and whole-cell lysate (WCL), and Actin (WCL) from stable adherent (SA) and suspended (SUSP) T24 cells. The graph represents ratio of densitometric band intensities as mean±SE from 3 independent experiments. **(H)** Western blot detection of Acetylated tubulin (Ace. tub) and Alpha tubulin (Alpha tub.) in whole-cell lysate from stable adherent (SA) and suspended (SUSP) T24 cells. The graph represents ratio of densitometric band intensities as mean±SE from 4 independent experiments. Scale bars-10 µm. Statistical analysis was done using Mann-Whitney’s test (non-normalized graphs), one-sample t-test (normalized graphs), and one-way Anova for the distribution profiles (* p-value<0.05, ** p-value <0.01, *** p-value <0.001, **** p-value <0.0001, ns = not significant).

### KIF5B as a possible regulator for cell-matrix adhesion-dependent Golgi organization

Cell-matrix adhesion regulates the activation of the small GTPase Arf1 (Singh et al., 2018) **(Fig. 3 D)**. Active Arf1 recruits minus-ended motor dynein to the Golgi, keeping it organized close to the microtubule-organizing center (MTOC) (Yadav et al., 2012; Singh et al., 2018). Upon loss of adhesion as the levels of active Arf1 drop, its binding to the Golgi membrane is expected to drop, causing dynein to be lost from the Golgi, promoting its disorganization (Singh et al., 2018). We asked if a plus-ended motor could also be involved in this pathway. The kinesin-1 family of motor proteins are reported to localize to the Golgi in multiple cell lines (Mahajan et al., 2019; Wobst et al., 2015). They also engage in a tug-of-war with dynein to regulate Golgi positioning in neuronal cells (Kelliher et al., 2018). Subcellular fractionation detects dynein and KIF5B in the Golgi-enriched membrane fraction **(Fig. 3 A)**. Immunostaining of KIF5B shows partial colocalization with ManII-GFP (cis-medial-Golgi marker) and GalT-RFP (trans-Golgi marker). Upon loss of adhesion, as active Arf1 is lost from the Golgi (B.R. et al., 2023), KIF5B localization also drops significantly from the cis and trans-Golgi. On early re-adhesion, as active Arf1 re-localizes to the Golgi (B.R. et al., 2023), KIF5B colocalization is also restored **(Fig. 3 B, C)**.

**Figure 3:**
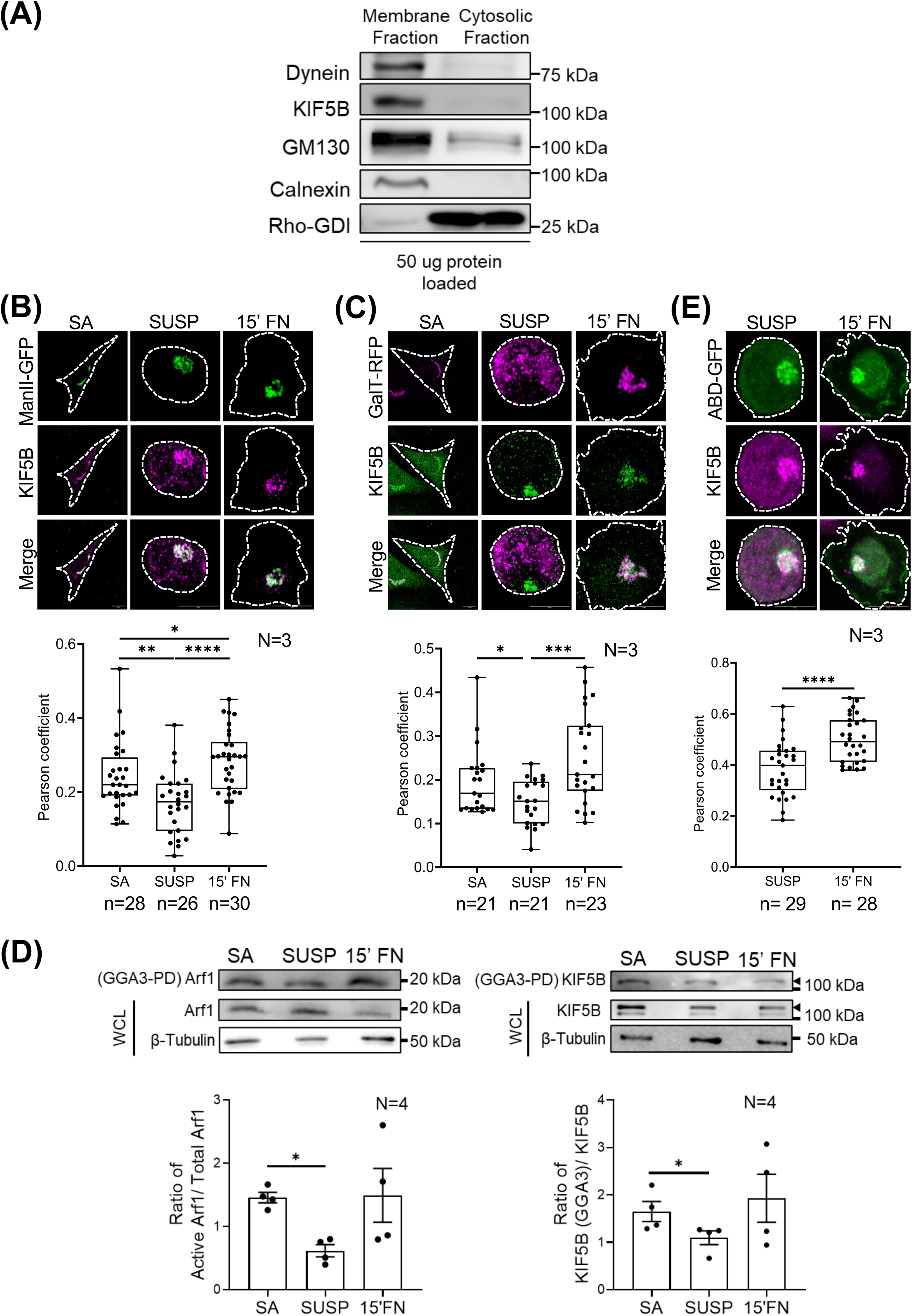
Role of Active Arf1 in recruiting Kinesin-1 to the Golgi: **(A)** Western blot detection of Dynein, KIF5B, GM130, Calnexin, and Rho-GDI in membrane and cytosolic fractions isolated from WT-MEFs. Equal protein was loaded. **(B, C)** Collapsed Z-stack images of stable adherent (SA), suspended (SUSP), and re-adherent (15′ FN) WT-MEFs, expressing **(B)** ManII-GFP and **(C)** GalT-RFP, immuno-stained for KIF5B. The graph shows Pearson colocalization coefficient of **(B)** ManII-GFP and **(C)** GalT-RFP with KIF5B using Z-stack images as median and quarters from 3 independent experiments. **(D)** Western blot detection of Arf1 (Left) and KIF5B (Right)(marked by ◄) in GST-GGA3 pulldown (GGA3 PD), whole-cell lysate (WCL), and β-Tubulin (WCL) from stable adherent (SA), suspended (SUSP), and re-adherent (15’ FN) WT-MEFs. The graph represents ratio of densitometric band intensities as mean±SE from 4 independent experiments. **(E)** Collapsed Z-stack images of suspended (SUSP) and re-adherent (15′ FN) WT-MEFs expressing ABD-GFP, immuno-stained for KIF5B. The graph shows the Pearson colocalization coefficient of ABD-GFP with KIF5B using Z-stack images as median and quarters from 3 independent experiments. Scale bars-10 µm. Statistical analysis was done using Mann-Whitney’s t-test (non-normalized graphs). (* p-value<0.05, ** p-value <0.01, *** p-value <0.001, **** p-value <0.0001, ns = not significant).

Active Arf1 has been shown to recruit KIF5B to lipid droplets (Rai et al., 2017), leading us to ask if it does the same at the Golgi in response to cell-matrix adhesion. GST-GGA3 pulldown of active Arf1 in stable adherent, suspended, and early re-adherent conditions show its levels drop upon loss of adhesion and recover upon early re-adhesion **(Fig. 3 D),** as reported earlier (Singh et al., 2018). KIF5B is detected in these active Arf1 pulldowns, and its binding decreases on the loss of adhesion as active Arf1 levels drop and recover upon re-adhesion **(Fig. 3 D)**. Further, KIF5B and active Arf1 (ABD-GFP), also colocalize at the Golgi in suspended and re-adherent cells, their overlap increasing in re-adherent cells **(Fig. 3 E)**. Further, expression of T31N-Arf1 (dominant negative Arf1) and the resulting breakup of the Golgi causes a reduction in its colocalization with KIF5B, as compared to WT-Arf1 and Q71L-Arf1 (constitutively active Arf1) expressing cells. This confirms the role active Arf1 has in mediating the recruitment of KIF5B to the Golgi **(Supp.** Fig. 3 **A)**. Alongside Arf1, different adaptor proteins could mediate the recruitment of dynein and KIF5B to the Golgi. This, in turn, could regulate Golgi organization and microtubule stability downstream of adhesion.

### Role of KIF5B and Dynein in regulating Golgi organization: Effect on cell function

To understand their relative roles in regulating Golgi organization, we used siRNA to deplete KIF5B and dynein, individually and together (dual KD) **(Fig. 4 A)**. The cis-Golgi marker, (GM130) revealed strikingly different phenotypes across these KDs. On KIF5B KD, the cis-Golgi disorganizes without being extensively dispersed. Dynein KD, however, causes the Golgi to disorganize and disperse throughout the cell. The dual KD of KIF5B and dynein prevents the Golgi from disorganizing and causes it to become compact, unlike their individual KDs **(Fig. 4 B, C)**. The percentage distribution and object count for these Golgi phenotypes are analyzed **(Fig. 4 B)**. Scrambled siRNA controls, for KIF5B and dynein, behaved similar to untreated control cells as seen using cis-Golgi marker (GM130) (**Supp. Fig. 4 A, B**). Cis-medial-Golgi marker Giantin also showed comparable phenotypes (**Supp. Fig. 4 C**). We further checked if pharmacological inhibition of dynein activity could produce Golgi phenotypes similar to dynein KD. Ciliobrevin D treatment inhibits the ATPase activity of dynein (Firestone et al., 2012), which is seen to cause Golgi dispersal similar to dynein KD, suggesting that functional endogenous protein is essential for Golgi organization **(Fig. 4 D, E)**. With no direct way to target the activity of KIF5B, we couldn’t test its impact on the Golgi organization. Ciliobrevin D treatment of KIF5B KD cells also causes the Golgi to have a compact phenotype similar to dual KD **(Fig. 4 D, E)**.

**Figure 4:**
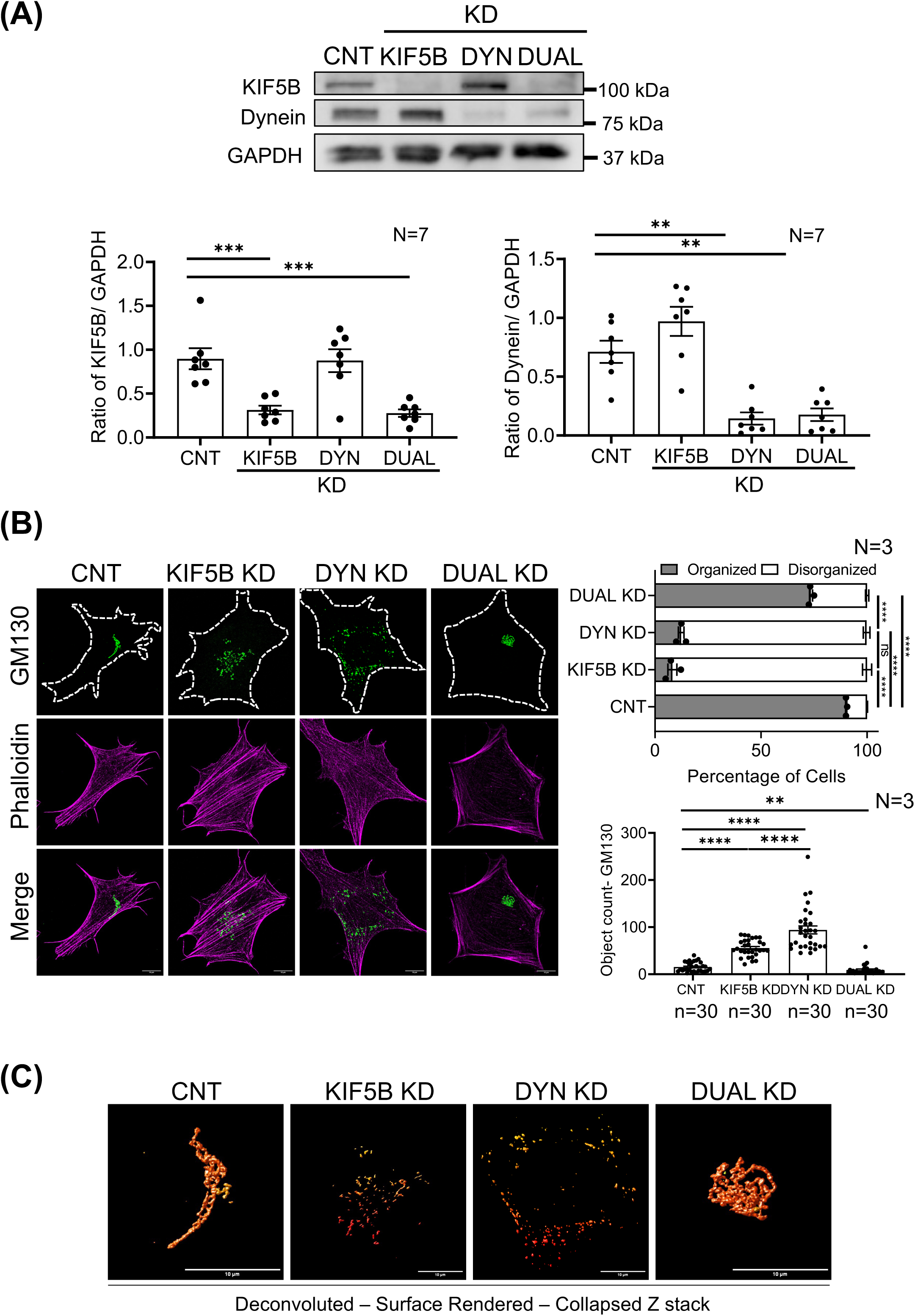
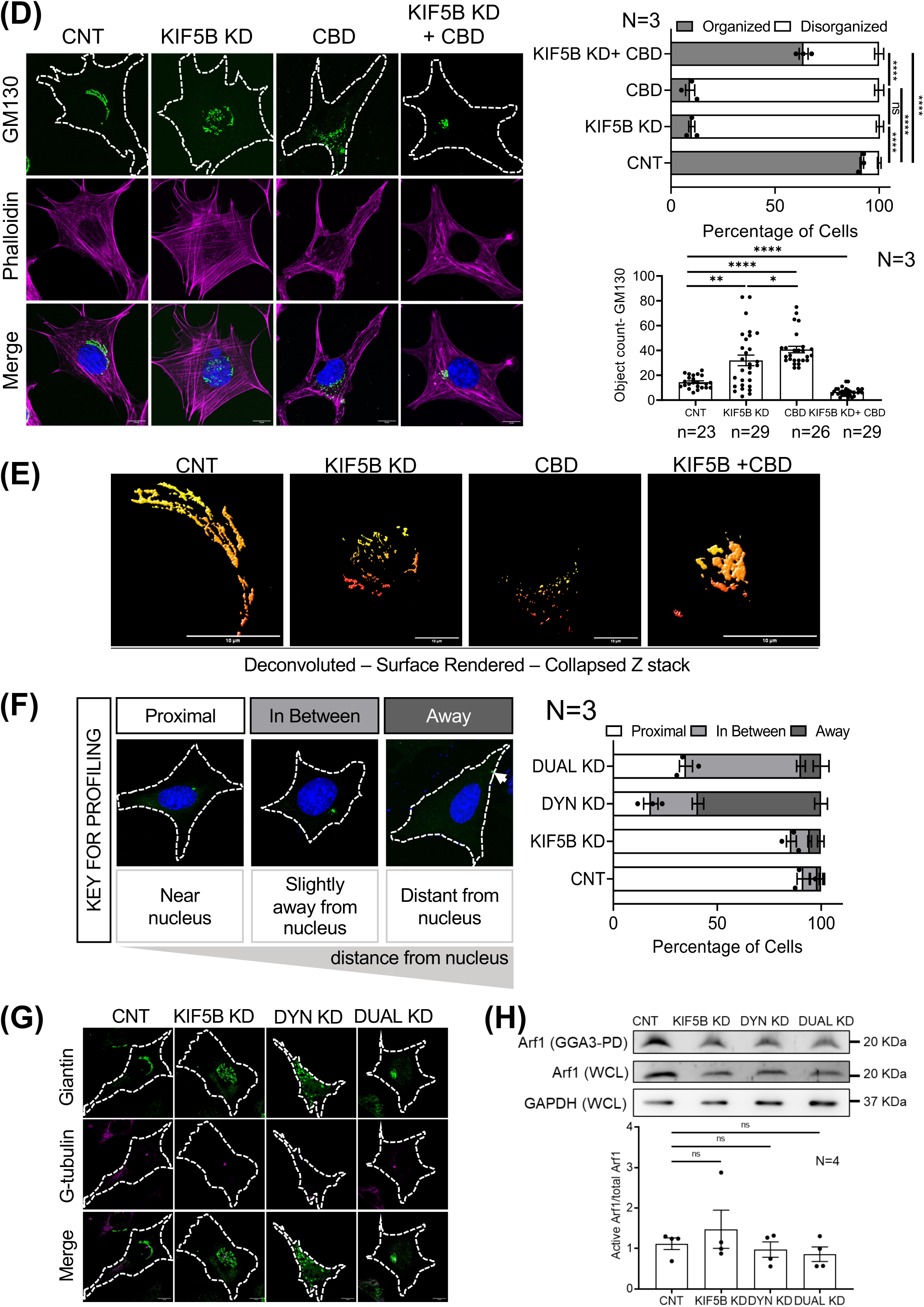
Role of KIF5B and Dynein in regulating Golgi organization and MTOC localization: **(A)** Western blot detection of KIF5B, dynein, and GAPDH from control WT-MEFs (CNT), or WT-MEFs treated with siRNA against KIF5B (KIF5B KD), dynein (DYN KD) individually, or together (DUAL KD). The graph represents ratio of densitometric band intensities as mean±SE from 7 independent experiments. **(B, C)** Representative deconvoluted **(B)** collapsed Z-stack and **(C)** surface-rendered collapsed Z-stack images of stable adherent control WT-MEFs (CNT), or WT-MEFs treated with siRNA against KIF5B (KIF5B KD), dynein (DYN KD) individually, or together (DUAL KD), immunostained with **(B)** GM130 and stained with Phalloidin, **(C)** GM130. In **(B),** the above graph shows the percentage distribution of cells with organized and disorganized Golgi phenotypes in these populations. The graph below represents discontinuous Golgi object count per cell. Both graphs are represented as mean±SE from 3 independent experiments. **(D, E)** Representative **(D)** cross-section and **(E)** deconvoluted surface rendered cross-section images of control WT-MEFs, or WT-MEFs treated with siRNA against KIF5B, treated with ciliobrevin (CBD) in control or KIF5B KD cells (KIF5B KD+CBD) immunostained with **(D)** GM130 and stained with DAPI and Phalloidin **(E)** GM130. The above graph in **(D)** shows the percentage distribution of cells with organized and disorganized Golgi phenotypes in these populations. The graph below represents discontinuous Golgi object count per cell. Both graphs are represented as mean±SE from 3 independent experiments. **(F)** Key for profiling shows cross-section images of WT-MEFs immunostained for Gamma-tubulin (green) and stained with DAPI (blue). The graph shows the percentage distribution of cells with MTOC localization concerning the nucleus as mean±SE from 3 independent experiments. **(G)** Collapsed Z-images of control WT-MEFs (CNT), or WT-MEFs treated with siRNA against KIF5B (KIF5B KD), dynein (DYN KD) individually, or together (DUAL KD), immunostained with Giantin and Gamma-tubulin (G-tubulin). **(H)** Western blot detection of Arf1 in GST-GGA3 pulldown (GGA3 PD), whole-cell lysate (WCL), and GAPDH (WCL) from control WT-MEFs (CNT), or WT-MEFs treated with siRNA against KIF5B (KIF5B KD), dynein (DYN KD) individually, or together (DUAL KD). The graph represents ratio of densitometric band intensities as mean±SE from 4 independent experiments. Scale bars-10 µm. Statistical analysis was done using Mann-Whitney’s test (non-normalized graphs) and one-way Anova for the distribution profiles (* p-value<0.05, ** p-value <0.01, *** p-value <0.001, **** p-value <0.0001, ns = not significant).

Loss of motor proteins has been reported to cause mislocalization of the centrosome (Splinter et al., 2010), which would affect the microtubule network and Golgi. KIF5B KD did not affect centrosome localization, while on loss of dynein, the centrosome is dramatically mislocalized to the cell periphery (away from the nucleus). Centrosome localization is partially restored on dual KIF5B and dynein KD **(Fig. 4 F)**. In control cells, the Golgi localizes near the centrosome while on KIF5B KD the disorganized Golgi is positioned around the centrosome as disconnected Golgi objects. In dynein KD cells, the Golgi is dispersed throughout the cell, with the centrosome positioned away from the nucleus. Interestingly, in dual KD cells, the Golgi localizes around the centrosome in a compact tubular organization **(Fig. 4 G)**. Further, we checked for and saw no change in active Arf1 levels in single KIF5B, dynein, and dual KD cells **(Fig. 4 H)**. This does raise the possibility that active Arf1 in KIF5B, dynein, and dual KD cells could contribute to the formation and localization of Golgi objects in these cells.

In dual KD cells, cis- and trans-Golgi are compact, with a juxtanuclear localization. The distinct Golgi ribbon organization seen in control cells is lost in the dual KD cells **(Fig. 5 A, B)**. This compact Golgi, continues to be linked to the microtubule network, as nocodazole treatment disperses the Golgi comparable to control cells **(Fig. 5 C)**. This also suggests that in the absence of KIF5B and dynein, other motor proteins, like KIFC3 from the kinesin-14 family, involved in Golgi positioning and integration (Xu et al., 2002) could keep the Golgi-microtubule association. It also reveals that KIF5B and dynein-mediated forces are likely needed to maintain the Golgi ribbon morphology and function.

**Figure 5:**
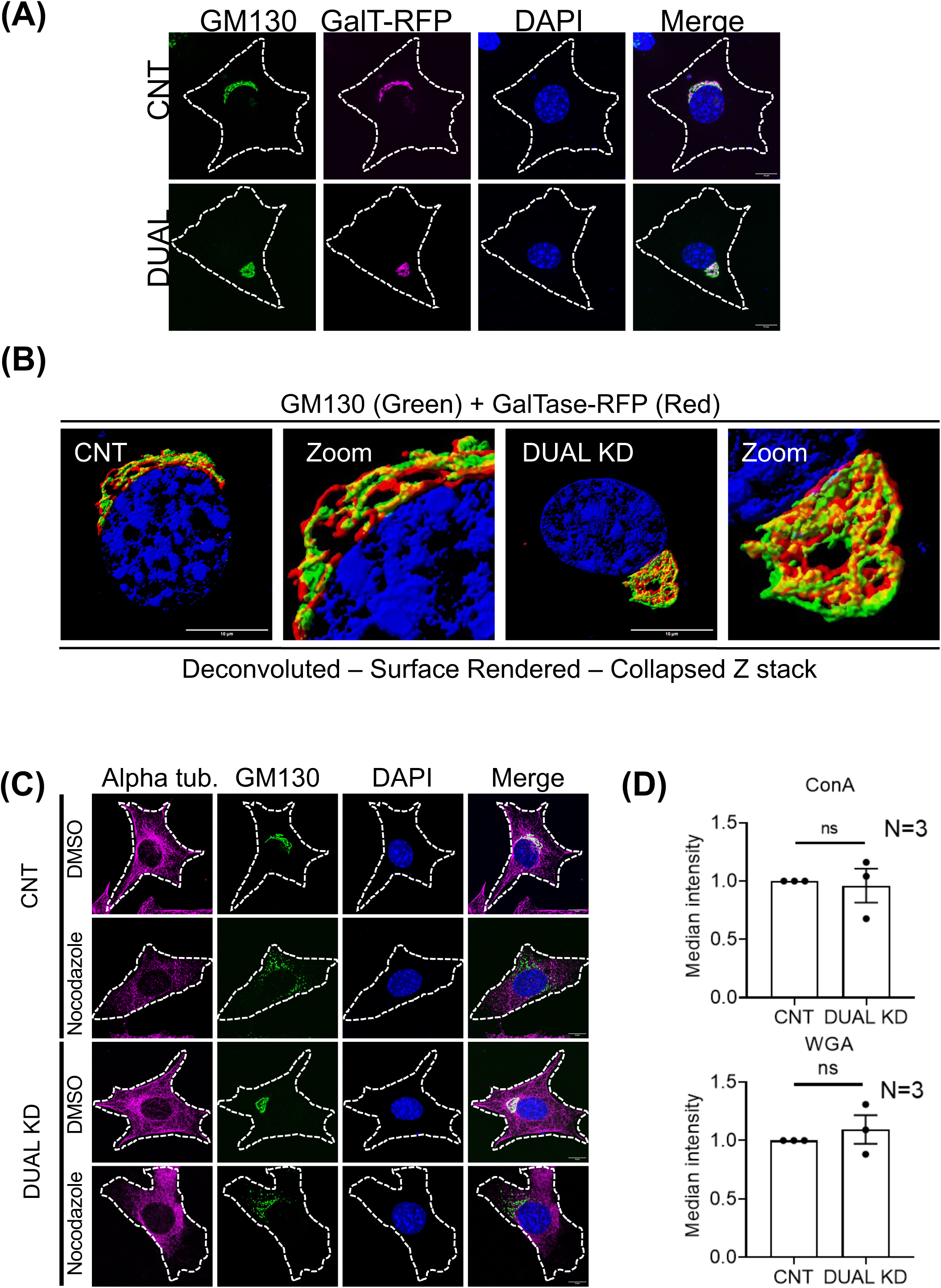
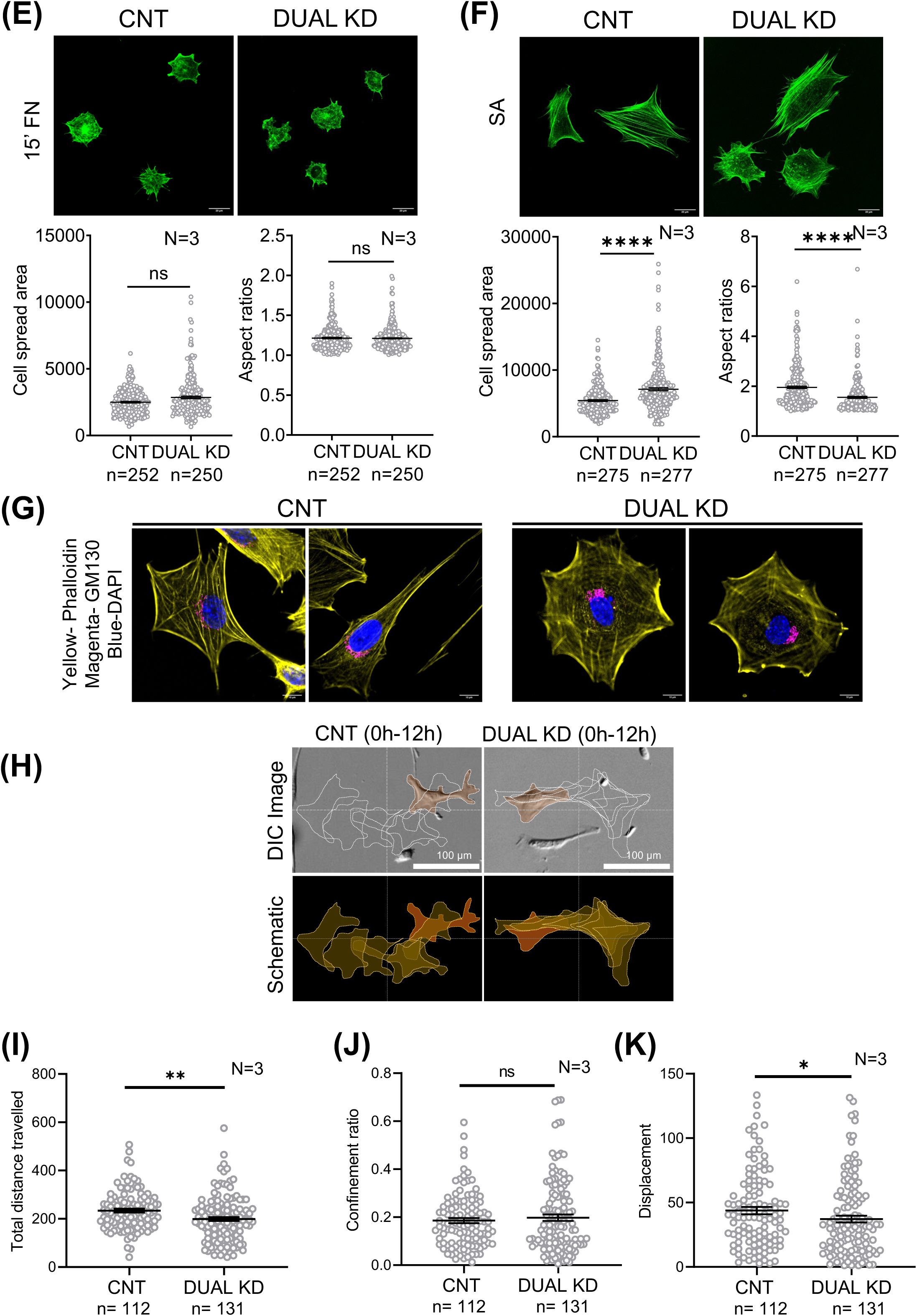
Golgi organization and cellular functions in KIF5B and Dynein dual knockdown cells: (A, **B)** Representative deconvoluted **(A)** collapsed Z-stack and **(B)** surface-rendered collapsed Z-stack images of stable adherent control WT-MEFs (CNT), or WT-MEFs treated with siRNA against KIF5B and dynein (DUAL KD), expressing GalT-RFP, immunostained with GM130, and stained with DAPI. **(C)** Collapsed Z-stack images of stable adherent WT-MEFs treated with DMSO or Nocodazole in control WT-MEFs (CNT), or WT-MEFs treated with siRNA against KIF5B and dynein (DUAL KD), immunostained for Alpha tubulin (Alpha tub.), GM130, and stained with DAPI. **(D)** Control WT-MEFs (CNT), or WT-MEFs treated with siRNA against KIF5B and dynein (DUAL KD), surface-labelled with ConA and WGA lectin. The graphs represent median fluorescence as mean±SE from 3 independent experiments. **(E, F)** Cross-section images of **(E)** re-adherent (15′ FN), and **(F)** stable adherent (SA) control WT-MEFs (CNT), or WT-MEFs treated with siRNA against KIF5B and dynein (DUAL KD), stained with phalloidin. The graphs represent the cell-spread area and aspect ratio as mean±SE from 3 independent experiments. **(G)** Cross-section images of stable adherent control WT-MEFs (CNT), or WT-MEFs treated with siRNA against KIF5B and dynein (DUAL KD), immunostained with GM130 and stained with phalloidin and DAPI. **(H)** Representative DIC image of WT-MEF treated with siRNA against KIF5B and dynein (DUAL KD), and their respective scrambled control siRNA (CNT), overlayed with tracing of the cell over 12 hours (using images at 2-hour intervals). The first image of the cell is highlighted in orange. The schematic below shows an overlay of the same cell tracings. **(I, J, K)** The graphs represent the quantitation of **(I)** total distance travelled, **(J)** displacement, and **(K)** confinement ratio of cells, as mean±SE from 3 independent experiments. Scale bars-10 µm, 20 µm, and 100 µm as indicated. Statistical analysis was done using Mann-Whitney’s test (non-normalized graphs) (* p-value<0.05, ** p-value <0.01, *** p-value <0.001, **** p-value <0.0001, ns = not significant).

A vital Golgi function in cells is their ability to glycosylate and deliver proteins and lipids to the plasma membrane. This process involves N- and O-glycosylation and depends on sequential enzyme activity across the cis, medial, and trans-Golgi compartments (Xiang et al., 2013; Pinho and Reis, 2015; Zhang and Wang, 2016). Changes in Golgi organization can alter glycoprotein processing and trafficking (Zhang and Wang, 2016; Pokrovskaya et al., 2011), which can be detected by lectin binding (Sharon, 2004). We evaluated changes in cell surface lectin binding in control vs. dual KD cells as a readout of Golgi function. Binding of fluorescently tagged ConA (mannose-binding) and WGA (galactose/ N-acetylgalactosamine-binding) were both unaffected in dual KD cells, relative to control cells **(Fig. 5 D)**. Changes in Golgi organization are also shown to affect cell polarity, and adhesion (Ravichandran et al., 2020; Ahat et al., 2019a). We evaluated the spreading and polarization of cells at early and late adhesion time points in control vs. dual KD cells. While no difference in both parameters was seen at early adhesion (15 minutes) **(Fig. 5 E),** in late adhesion cells, the dual KD cells spread more and became less polar than control cells **(Fig. 5 F)**. The Golgi is expectedly positioned at the front of a migrating cell (Ravichandran et al., 2020) **(Fig. 5 G)**, which is affected in dual KD cells **(Fig. 5 G)**. This could contribute to how the microtubule network is arranged in cells, affecting directional vesicle trafficking, polarity, and cell migration (Small and Kaverina, 2003; Ravichandran et al., 2020). Single cell migration analysis shows that upon dual KD, total distance and net displacement were both less compared to control **(Fig. 5 H, I, J)**. However, a comparable confinement ratio **(Fig. 5 K)** suggests directional migration of dual KD cells is not significantly affected. The compact Golgi in dual KDs could affect the speed and displacement seen, but can regulate directional migration via regulation of microtubule network organization (Kopf and Kiermaier, 2021; Garcin and Straube, 2019).

### KIF5B and Dynein-mediated Golgi organization: Effect on microtubule network organization

Golgi is seen to have a role in regulating microtubule nucleation and stabilization (Ryan et al., 2012), as previously discussed. Loss of adhesion-mediated Golgi disorganization affects microtubule acetylation and detyrosination **(Fig. 1 A, B, C)**. This led us to ask if motor protein-dependent changes in Golgi organization differentially affect microtubule stability. Western blot studies show microtubule acetylation levels not to be significantly affected by motor protein KD (single and dual) **(Fig. 6 A)**. We further asked if microtubule network organization could be affected. Acetylated tubulin localized near the Golgi in stable adherent cells **(Fig. 1 D)** is dispersed as the Golgi disorganizes in KIF5B and dynein KD cells **(Fig. 6 B)**. In dual KD cells, the compact Golgi partially restores the acetylated tubulin network **(Fig. 6 B)**. The alpha tubulin network is unaffected across all single and dual KDs **(Supp. Fig. 6 A)**. An analysis pipeline developed for mitochondrial morphology (Chaudhry et al., 2020) was adapted for assessing tubulin network organization. Branch number analysis in Z-stack images reflects the density of microtubules. This reveals acetylated tubulin branch numbers to drop without affecting branch length in KIF5B KD and dynein KD cells (**Fig. 6 C, D)**. Branch numbers are interestingly restored in dual KD cells, suggesting their compact Golgi could regulate acetylated microtubule organization in cells **(Fig. 6 D)**. Alpha tubulin branch number and branch length were unaffected by single KD or dual KD **(Fig. 6 E, F)**.

**Figure 6:**
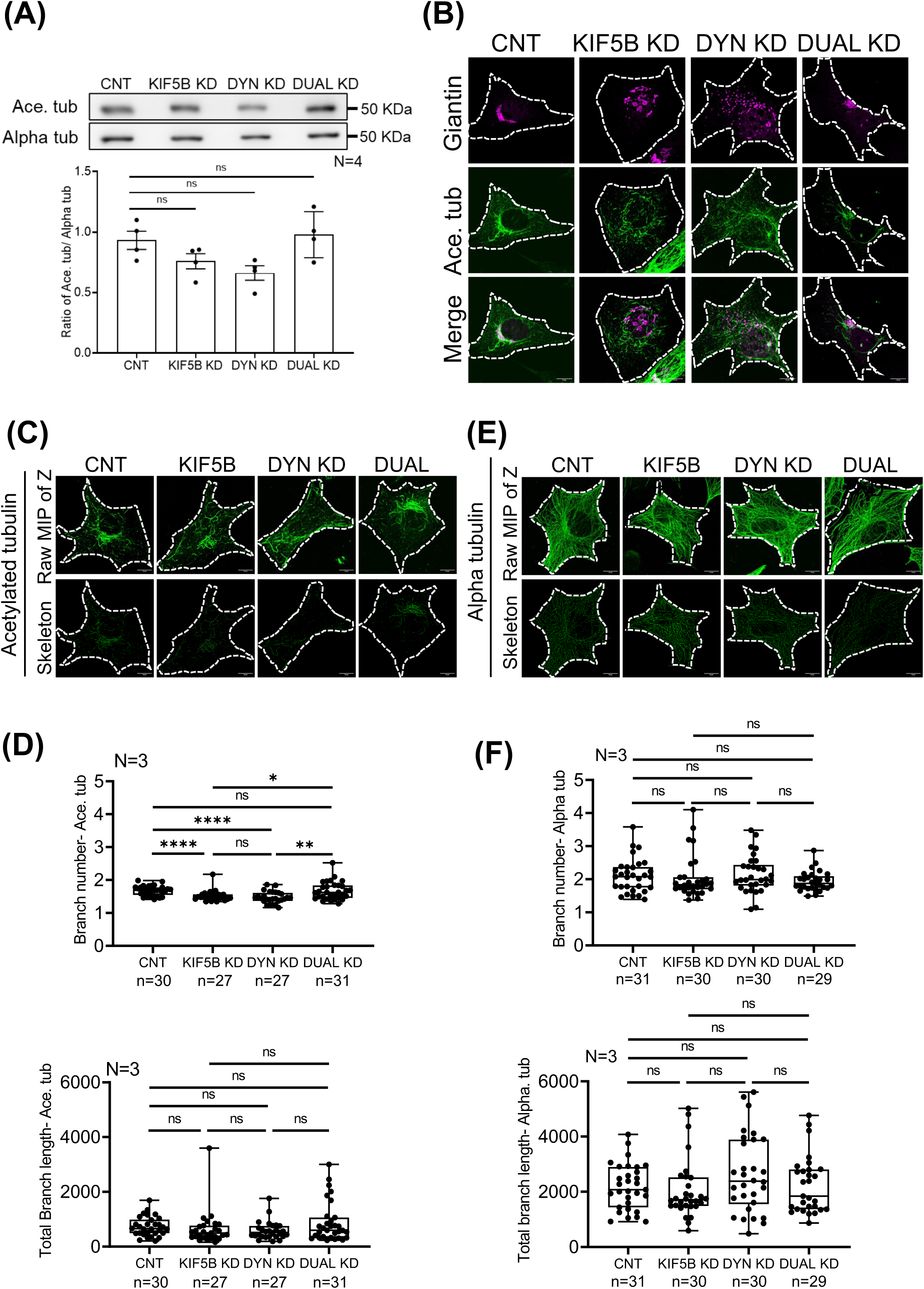
Role of KIF5B and Dynein in Golgi organization mediated regulation of MT architecture: **(A)** Western blot detection of Acetylated tubulin (Ace. tub) and Alpha tubulin (Alpha tub) in whole-cell lysate from control WT-MEFs (CNT), or WT-MEFs treated with siRNA against KIF5B (KIF5B KD), dynein (DYN KD) individually, or together (DUAL KD). The graph represents ratio of densitometric band intensities as mean±SE from 4 independent experiments. **(B, C, E)** Collapsed Z-stack images of stable adherent control WT-MEFs (CNT), or WT-MEFs treated with siRNA against KIF5B (KIF5B KD), dynein (DYN KD) individually, or together (DUAL KD), immunostained with **(B)** Giantin and Acetylated tubulin (Ace. tub) or **(C)** Acetylated tubulin (Ace. tub) and **(E)** Alpha tubulin (Alpha tub.). **(D, F)** The graphs represent the quantitation of branch number and total branch length as median and quarters for **(D)** Acetylated-tubulin (Ace. tub) and **(F)** Alpha tubulin (Alpha tub.) from 3 independent experiments. Scale bars-10 µm. Statistical analysis was done using Mann-Whitney’s test (non-normalized graphs) (* p-value<0.05, ** p-value <0.01, *** p-value <0.001, **** p-value <0.0001, ns = not significant).

### KIF5B and Dynein regulate Golgi organization on loss of adhesion and re-adhesion

When adhesion is lost, the Golgi is distinctly disorganized (Singh et al., 2018), which could be mediated by the differential role and regulation of dynein and KIF5B. This could further depend on the adhesion-dependent active Arf1 that can bind KIF5B and dynein, possibly recruiting them to the Golgi. Upon loss of adhesion, as active Arf1 levels drop **(Fig. 3 D)**, its binding to KIF5B **(Fig. 3 D)** and dynein decreases (Singh et al., 2018). The relative loss of KIF5B vs. dynein on loss of adhesion could help reveal their relative role in Golgi disorganization. KIF5B KD causes the Golgi to disorganize significantly, while in dynein KD cells, the Golgi is completely dispersed. In dual KD cells, the Golgi stays compact on loss of adhesion **(Fig. 7 A),** failing to disorganize. This suggests KIF5B and dynein at the Golgi are vital for the observed Golgi disorganization phenotype on loss of adhesion. Early re-adhesion of these suspended cells causes the Golgi to reorganize rapidly in control cells and KIF5B KD cells **(Fig. 7 B)**. In dynein KD cells, the Golgi fails to reorganize **(Fig. 7 B)**, possibly reflecting the disruption of MTOC localization **(Fig. 4 F)**, as is also seen on Ciliobrevin D treatment (Singh et al., 2018). Dual KD cells keep their compact Golgi phenotype on early re-adhesion as well **(Fig. 7 B)**.

**Figure 7:**
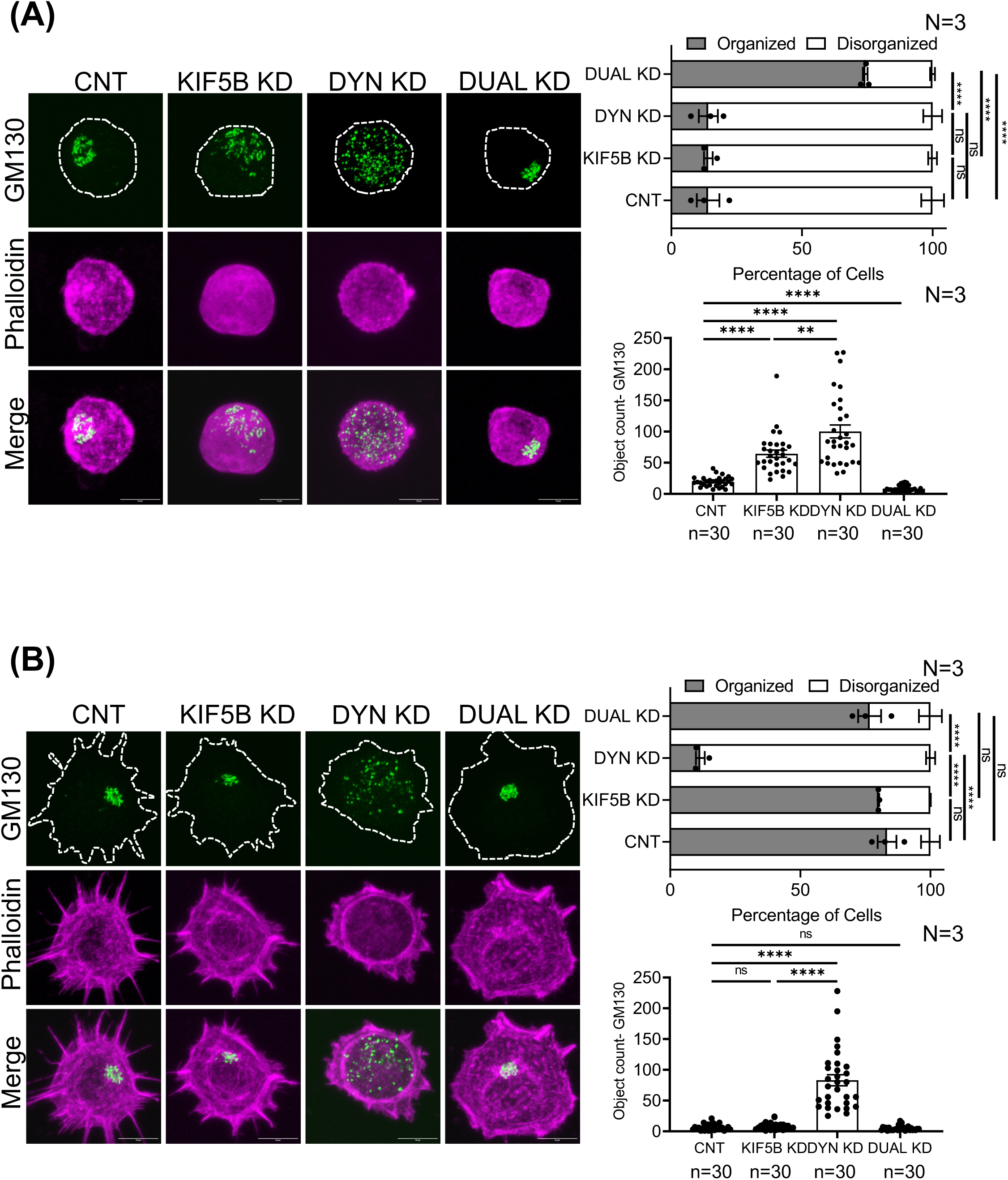
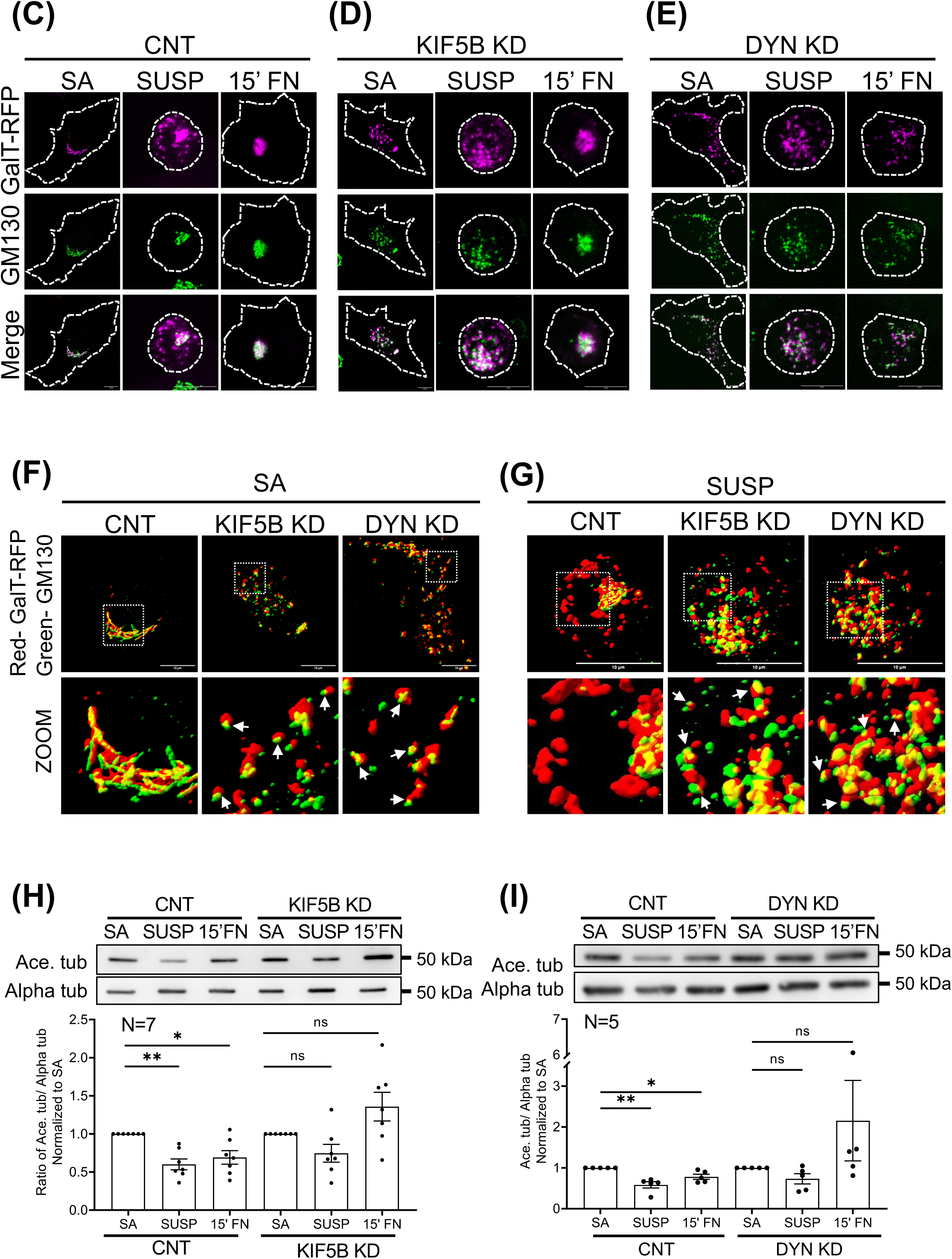
Role of KIF5B and Dynein in regulating Golgi organization upon loss of adhesion and early re-adhesion- Role in regulating MT Stability: (A, **B)** Collapsed Z-stack images of **(A)** suspended and **(B)** early re-adherent control WT-MEFs (CNT), or WT-MEFs treated with siRNA against KIF5B (KIF5B KD), dynein (DYN KD) individually, or together (DUAL KD), immunostained with GM130 and stained with Phalloidin. The above graph shows the percentage distribution of cells with organized and disorganized Golgi phenotypes in these populations. The graph below represents discontinuous Golgi object count per cell. Both graphs are represented as mean±SE from 3 independent experiments. **(C, D, E)** Collapsed Z-stack images of **(C)** control (CNT), **(D)** KIF5B, and **(E)** dynein (DYN) KD stable adherent (SA), suspended (SUSP), and re-adherent (15′ FN) WT-MEFs, expressing GalT-RFP and immunostained with GM130. **(F, G)** Collapsed Z-stack and deconvoluted surface rendered images of **(F)** stable adherent (SA) and **(G)** suspended (SUSP) WT-MEFs, expressing GalT-RFP and immunostained with GM130. Golgi ministacks are marked with arrows. **(H, I)** Western blot detection of Acetylated (Ace. tub) and Alpha tubulin (Alpha tub) in whole-cell lysate from control WT-MEFs (CNT) and WT-MEFs treated with siRNA against **(H)** KIF5B (KIF5B KD), and **(I)** dynein (DYN KD). The graph represents ratio of densitometric band intensities as mean±SE from 7 and 5 independent experiments, respectively. Scale bars-10 µm. Statistical analysis was done using Mann-Whitney’s test (non-normalized graphs), one-sample t-test (normalized graphs), and one-way Anova for the distribution profiles (* p-value<0.05, ** p-value <0.01, *** p-value <0.001, **** p-value <0.0001, ns = not significant).

As previously discussed, cis and trans-Golgi are essential for regulating microtubule nucleation and stabilization. On loss of adhesion the differential disorganization of the cis/cis-medial-Golgi from the trans-Golgi **(Fig. 7 C)** is unique and vital to Golgi function. We hence evaluated the relative organization of cis- and trans-Golgi in suspended and re-adherent KIF5B and dynein KD cells. Staining for cis-Golgi marker GM130 in trans-Golgi marker GalT-RFP expressing cells shows KIF5B **(Fig. 7 D, F and G),** and dynein KD **(Fig. 7 E, F and G)** to form distinct Golgi ministacks, where cis- and trans-Golgi localized together in adherent **(Fig. 7 F)** and suspended **(Fig. 7 G)** cells. Such Golgi ministacks are known to form upon Jaw1/LRMP KD, regulating acetylated tubulin levels, but not localization (Okumura et al., 2023). In KIF5B and dynein KD cells, Golgi ministacks in stable adherent cells **(Fig. 7 F)** maintain microtubule acetylation levels **(Fig. 6 A)**; but not their organization **(Fig. 6 C, D)**. Interestingly, the presence of Golgi ministacks regulates microtubule acetylation levels also upon loss of adhesion, as detected by western blot **(Fig. 7 H and 7 I)**. Together, they not only suggest a role for KIF5D and dynein in regulating adhesion-dependent Golgi organization, but also reveal how disrupting these could lead to the formation of differential Golgi ministacks and a compact Golgi that are functionally distinct.

## DISCUSSION

Cell-matrix adhesion-dependent regulation of cellular processes are mediated by their control of cell signalling and cytoskeletal organization (Bershadsky et al., 1996; Buwa et al., 2021; Small and Kaverina, 2003; Tran et al., 2007; Gaus et al., 2006). Regulation of actin, microtubule networks, and their respective motors relays important adhesion-dependent signalling cues to organelles, regulating their organization, localization, and function (Bershadsky et al., 1996; Copeland et al., 2016). Several motor protein families of kinesin, dynein, myosins, and non-conventional GTPase dynamin are associated with the Golgi complex, regulating Golgi morphology and function (Horgan et al., 2010; Lee et al.; She et al., 2017; Miserey-Lenkei et al., 2017). Adhesion-dependent regulation of Arf1 activation and its dynein binding recruits it to the Golgi, regulating its organization and function, like cell surface glycosylation (Singh et al., 2018). This study shows a possible role for Arf1 in recruiting KIF5B alongside dynein, to Golgi, highlighting the critical role that plus and minus-ended motor protein forces have in Golgi organization. Upon loss of adhesion, differential loss of active Arf1, plus and minus-ended motors, could drive differential disorganization of the cis and trans-Golgi (Singh et al., 2018), further regulating trafficking, protein, and lipid processing (Pulvirenti et al., 2008; Glick and Luini, 2011).

In mammalian cells, connected Golgi stacks form unique ribbon-like structures contributing to higher-order Golgi architecture (Benvenuto et al., 2024). This ribbon-like organization can vary between cell types and be actively regulated. Assembled using an intact microtubule network, associated motors ensure proper positioning and organization of the Golgi (Copeland et al., 2016; Kelliher et al., 2018; Yadav et al., 2012). The LINC (linker of nucleoskeleton and cytoskeleton) complex (containing SUN2/nesprin2), along with KIF20A, regulates the juxtanuclear organization of the Golgi (Hieda et al., 2021). Further, nesprins recruit kinesin-1 to the nuclear membrane (Wilson and Holzbaur, 2015), which could further regulate Golgi positioning. The microtubule-Golgi crosstalk is now known to be bidirectional, with the Golgi housing vital microtubule nucleating and stabilizing factors, like AKAP450 (Rivero et al., 2009). Trans-Golgi and TGN localized CLASPs are microtubule-binding proteins that selectively coat non-centrosomal microtubule seeds, which regulate asymmetric Golgi-derived microtubule network (Miller et al., 2009; Efimov et al., 2007). Golgi-derived preferentially acetylated microtubules support directional post-Golgi traffic, vital for cell polarization and directional cell migration (Miller et al., 2009; Efimov et al., 2007). Although what causes preferential acetylation of the Golgi-derived microtubules remains unclear, studies suggest that distinct structural properties of these tubules could impact the accessibility of ATAT-1 and HDAC6/SIRT2 (Donker and Godinho, 2025). Adhesion scaffold proteins like paxillin (Deakin and Turner, 2014), and matrix stiffness cues also regulate microtubule acetylation (Wen et al., 2023). Golgi is a mechanoresponsive organelle (Théry and Pende, 2019), its organization is affected by matrix stiffness, possibly downstream of adhesion (Saha et al., 2025).

Our study shows an adhesion-dependent drop and recovery of microtubule acetylation and detyrosination upon loss of adhesion and re-adhesion to be independent of total paxillin levels that remain unchanged. The close association between Golgi organization and acetylated tubulin suggests a possible cause-and-effect relationship between adhesion-dependent Golgi organization and microtubule stability, which we show are closely regulated. Upon loss of adhesion as Arf1 activity drops, the dissociation of cis and trans-Golgi could affect the ability of the Golgi to nucleate and/or stabilize microtubules, affecting tubulin acetylation levels. While active Arf1 expression (Q71L-Arf1) in non-adherent cells restores Golgi organization and microtubule acetylation, a small but significant increase in microtubule acetylation is also observed in Q71L-Arf1 expressing cells. This suggests that both Golgi organization and adhesion-dependent Arf1 activation could contribute to microtubule acetylation levels. In T24 bladder cancer cells, anchorage-independent Arf1 activation supports Golgi organization, keeping the cis- and trans-Golgi together. This allows T24 cells to keep microtubule acetylation levels upon loss of adhesion, which may affect cell function. The impact of the differential expression of motor proteins in cancers on Golgi organization and microtubule acetylation becomes of particular interest. Like the expression of Arf1 (Xie et al., 2016), kinesin levels are variable in cancers (Moamer et al., 2019), and can even be potential pancreatic cancer biomarker (Charles Jacob et al., 2022).

A kinesin-dynein tug-of-war regulates Golgi positioning in neuronal cells (Kelliher et al., 2018). Active Arf1 could fine-tune the KIF5B and dynein association with the Golgi, and a possible tug-of-war to regulate Golgi disorganization on loss of adhesion and re-organization upon re-adhesion (Kelliher et al., 2018). Different adaptors like bicaudal D2 (Splinter et al., 2010), HOOK3 (Walenta et al., 2001; Cross and Dodding, 2019), TRAK2, RILP (Cross and Dodding, 2019), etc, bind opposing motors to regulate organelle positioning. They could also be co-recruiting KIF5B and dynein to the Golgi in association with Arf1 in adherent and non-adherent cells. siRNA-mediated KD of KIF5B and dynein disrupts Golgi organization, causing them to form distinct ministacks **(Fig. 8)**, as functional Golgi subunits (Copeland et al., 2016). Active Arf1 levels maintained in the KIF5B and dynein KD cells, could continue recruiting motor protein to Golgi ministacks **(Fig. 8)**. In KIF5B KD cells, ministacks localize around the nucleus and MTOC, while in dynein KD and upon pharmacological inhibition of dynein, Golgi ministacks are dispersed throughout the cell. This suggests that dynein could support Golgi ministack localization in KIF5B KD cells. KIF5B and other kinesins associated with the Golgi could similarly influence the localization of Golgi ministacks in dynein-targeted cells **(Fig. 8)**. Knowing Arf1 activation is unaffected by the loss of KIF5B or dynein, Arf1 localization to Golgi mini stacks could have implications for their positioning and function in cells. As reported earlier in T24 cells, anchorage-independent Arf1 activation could also impact the formation of Golgi ministacks in cancer cells.

**Figure 8:**
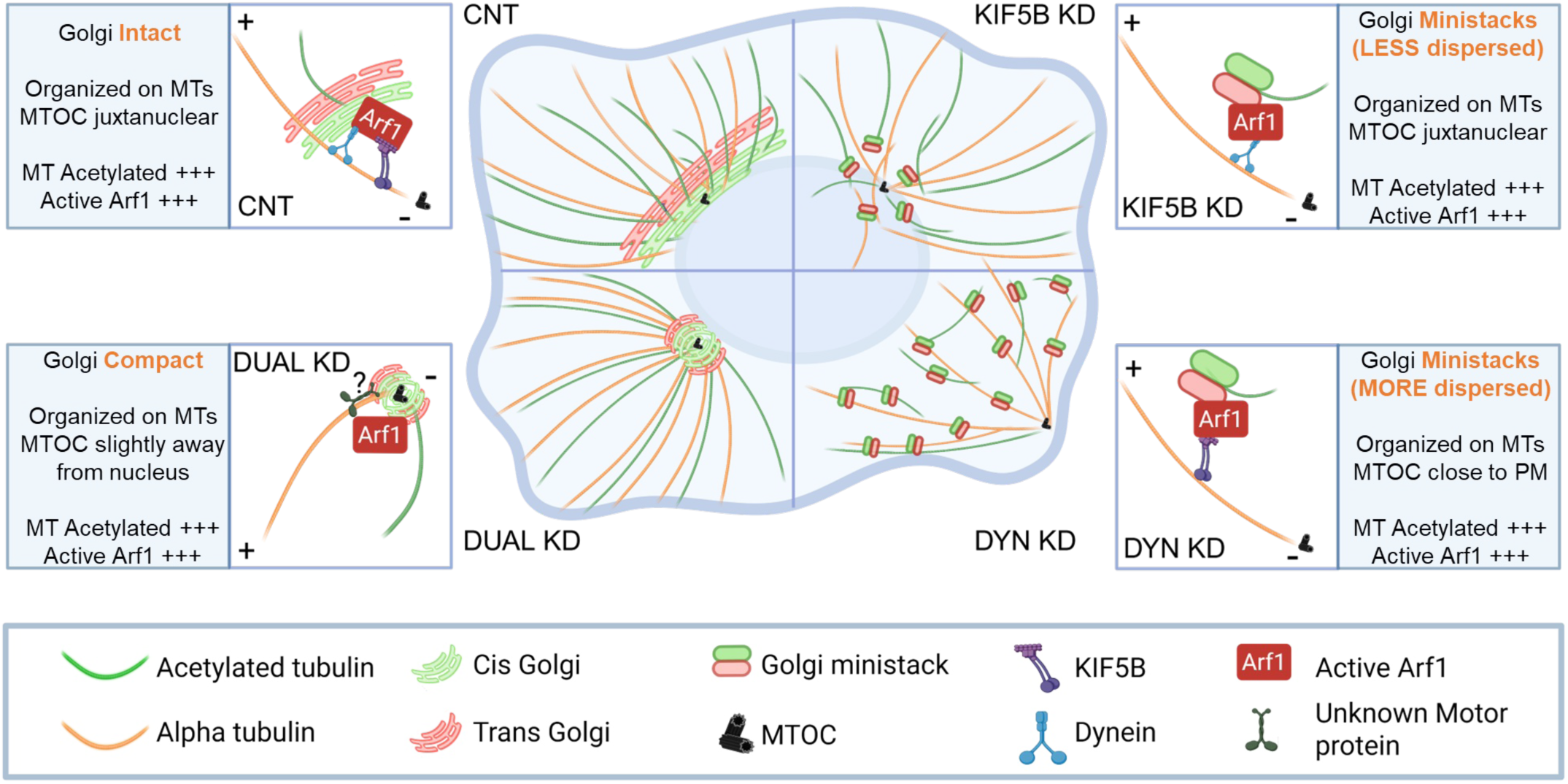
KIF5B and dynein regulate Golgi organization and localization: Role in regulating microtubule acetylation: In control (CNT) cells, active Arf1 facilitates the recruitment of KIF5B and dynein to the Golgi, maintaining its juxtanuclear organization. Knockdown of either KIF5B (KIF5B KD) or dynein (DYN KD) disrupts Golgi organization, forming Golgi ministacks with cis- and trans-Golgi. In KIF5B KD cells, active Arf1 could recruit dynein to ministacks, positioning them near the microtubule-organizing center (MTOC). In DYN KD cells, MTOC is localized close to the plasma membrane. This combined with active Arf1-recruited KIF5B causes the dispersal of ministacks. In cells joint knockdown of both motors (DUAL KD), causes the Golgi to become compact around the MTOC, keeping their cis- and trans-Golgi localization. Across all conditions, the Golgi remains microtubule-associated and supports microtubule acetylation, similar to control cells.

This also raises a query about how the loss of adhesion-mediated regulation of Arf1 activity could control relative levels of dynein and KIF5B at the Golgi to support their distinctly disorganized phenotype. It would be of interest to determine whether the differential levels of active Arf1 in the cis-/cis-medial vs. trans-Golgi contribute to the differential dynein and KIF5B recruitment to regulate their spatial distribution in cells. The implications of such a recruitment in maintaining Golgi ministacks are also worth evaluating, as is the role of other motor proteins.

Golgi ministacks can regulate microtubule acetylation levels (Okumura et al., 2023). Nascent tubules are nucleated from the cis-face of the Golgi (Rivero et al., 2009), which are then stabilized by GRASP65 (Ayala et al., 2019). Golgi ministacks in KIF5B and dynein KD cells maintain the tubulin acetylation levels, however, the organization of acetylated tubulin network is disrupted, as reflected from the drop in tubule density (branch number). In dual KD, the compact Golgi, unlike ministacks, restores acetylated microtubule branching. Interestingly, on KIF5B and dynein KD, Golgi ministacks prevent the loss of adhesion-mediated drop in microtubule acetylation. These functional Golgi outcomes, retaining on loss of adhesion, suggests KIF5B and dynein constitute major adhesion-dependent regulators contributing to loss-of-adhesion Golgi phenotypes. This also indicates an important role for cell-matrix adhesion and its regulation of microtubule acetylation via Golgi organization. Such Golgi organization-mediated regulation of microtubule acetylation in cancer cells could further aid cell proliferation, migration, and metastasis (Donker and Godinho, 2025; Deakin and Turner, 2014; Bance et al., 2019; Wen et al., 2023). On re-adhesion, the re-organization of the Golgi in KIF5B KD cells suggests dynein to be vital for adhesion-dependent regulation of the Golgi, as has also been reported earlier (Singh et al., 2018).

Dynein KD also dramatically mislocalizes the MTOC to the cell periphery, as seen in our studies and reported earlier in U2OS cells (Splinter et al., 2010). During Jurkat T cell activation, Cytoplasmic linker protein 170 (CLIP70) phosphorylation regulates the plus end-tracking of dynein, which repositions MTOC from the cell periphery to the juxtanuclear region (Lim et al., 2018). MTOC positioning is vital for microtubule network organization, which regulates organelles’ localization and organization. Directional cargo trafficking, also governed by the microtubule network organization (Balabanian et al., 2018), regulates different cellular processes (Millarte and Farhan, 2012; Kaverina and Straube, 2011; Bance et al., 2019).

The dual KD of KIF5B and dynein, causing the Golgi to stay compact around the MTOC, suggests their joint presence could have a role in keeping the classical Golgi ribbon morphology. A compact Golgi phenotype around the MTOC was earlier reported in Microtubule crosslinking factor 1 (MTCL-1) KD cells. MTCL-1 mediates juxtanuclear accumulation of stable microtubules and their crosslinking, which promotes Golgi microtubule linkage and supports Golgi ribbon formation (Sato et al., 2014). Loss of KIF5B and dynein affects the balancing forces on the Golgi, causing its localization at the MTOC to be retained. This holds on loss of adhesion and early re-adhesion. In some ways, it strengthens our suggestion that KIF5B and dynein could be the significant adhesion-dependent Golgi regulators at play. The dual KD compact Golgi, in staying associated with microtubules, suggests a plausible role for minus-ended kinesins like KIFC3 (Xu et al., 2002) and plus-ended KIF1C (Lee et al.), Eg-5 (Wakana et al., 2013), kinesin-2 (Stauber et al., 2006), and KIF25 (Liu et al., 2019), in Golgi organization. The fact that these kinesins are implicated in Golgi positioning and Golgi-derived traffic further strengthens their possible involvement (Xu et al., 2002; Lee et al.; Stauber et al., 2006; Wakana et al., 2013; Liu et al., 2019).

The compact Golgi in dual KD cells also keeps their relative localization of cis- and trans-Golgi, not affecting early adhesion-dependent cell spreading and cell aspect ratio (polarity). In late adherent dual KD cells, a distinct increase in cell spreading, accompanied by a decrease in polarity, suggests that the compact Golgi could indeed affect cell behaviour. The differentially localized Golgi and its organization could impact directional trafficking and processing to affect cell migration (Ahat et al., 2019b; Ebnet et al., 2018; Yadav and Linstedt, 2011; Linstedt, 2004). This produces a distinct effect on speed and displacement of dual KD cells, without affecting their overall directionality of migration. This could be the result of the compact Golgi keeping its juxtanuclear localization, which can regulate the microtubule network in these cells. This also suggests the characteristic juxtanuclear ribbon Golgi morphology in cells maintained by the presence of KIF5B and dynein is vital to Golgi and possibly cellular functions.

PTMs of microtubules in working with microtubule-associated proteins (MAPs) can modulate the binding and motility of specific motors (Wloga and Gaertig, 2010), which in turn regulate directional cargo traffic. Acetylation of tubulin promotes the binding of kinesin-1 and dynein (Dunn et al., 2008; Cai et al., 2009; Dompierre et al., 2007). The selective interaction of kinesin-1 with acetylated microtubules is likely mediated by specific MAPs, which may modulate motor-microtubule affinity in an acetylation-dependent manner (Andreu-Carbó et al., 2024). MAP7 increases the binding rate of kinesin-1 to microtubules, regulating organelle plus-end targeting (Chaudhary et al., 2019). Depletion of Hsp90 (known to bind MAP7) leads to a decrease in microtubule acetylation, leading to Golgi fragmentation and impaired anterograde traffic (Wu et al., 2020). Understanding the MAP and PTM landscapes upon loss of adhesion and changing matrix stiffness could give physiological insights into how these could affect the binding and motility of motors along the microtubule network, influencing Golgi organization and function. The differential regulation of Arf1, Golgi organization, and microtubule acetylation in T24 bladder cancer cells suggests this regulatory crosstalk could further support anchorage-independent signalling in cancers.

## Acknowledgement

A SERB CRG grant supports this work – CRG/2022/001813 to NB. AC, RBR and NBu are supported by a fellowship from the Council of Scientific & Industrial Research (CSIR). We acknowledge the extensive support the IISER Pune Microscopy Facility provides for cellular imaging and the IISER Pune FACS Facility.

## Author Contributions

The experimental work was led by AC. Data was recorded, organized, and analyzed by AC, SP, RBR, NBu, and AD. Data Analysis for some experiments were done by RB and MJ. Analysis and representation of all data has support from NB. The manuscript was written by AC and NB. All authors have read and given approval to the final version of the manuscript. The authors declare no competing financial interest.

---

**Figure 1 Supplementary:** Western blot detection of Paxillin and GAPDH in the whole-cell lysate from stable adherent (SA), suspended (SUSP), and re-adherent (15′ FN) WT-MEFs. The graph represents ratio of densitometric band intensities as mean±SE from 4 independent experiments. Statistical analysis was done using Mann-Whitney’s t-test (non-normalized graphs) (* p-value<0.05, ** p-value <0.01, *** p-value <0.001, **** p-value <0.0001, ns = not significant).

**Figure 3 Supplementary:** Collapsed Z-stack images of stable adherent WT-MEFs expressing WT-Arf1, T31N-Arf1, and Q71L-Arf1, immuno-stained for KIF5B and GM130. The graph shows the Pearson colocalization coefficient of GM130 with KIF5B using Z-stack images as median and quarters from 3 independent experiments. Scale bars-10 µm. Statistical analysis was done using Mann-Whitney’s test (non-normalized graphs). (* p-value<0.05, ** p-value <0.01, *** p-value <0.001,**** p-value <0.0001, ns = not significant).

**Figure 4 Supplementary: (A)** Western blot detection of KIF5B (marked by ◄), dynein, and GAPDH from control WT-MEFs (CNT), or WT-MEF treated with siRNA individually against KIF5B (KIF5B KD) and dynein (DYN KD), or together (DUAL KD), and their respective scrambled control siRNA (Scr Dual). **(B)** Collapsed Z-stack images of stable adherent control WT-MEFs (CNT), or WT-MEF treated with siRNA individually against KIF5B (KIF5B KD) and dynein (DYN KD), or together (DUAL KD), and their respective scrambled control siRNA (Scr Dual), immunostained with GM130 and stained with Phalloidin. The above graph shows the percentage distribution of cells with organized and disorganized Golgi phenotypes in these populations. The graph below represents discontinuous Golgi object count per cell. Both graphs are represented as mean±SE from 3 independent experiments. **(C)** Collapsed Z-stack images of stable adherent control WT-MEFs (CNT), or WT-MEF treated with siRNA individually against KIF5B (KIF5B KD) and dynein (DYN KD), or together (DUAL KD), immunostained with Giantin. The graph represents discontinuous Golgi object count per cell mean±SE from 3 independent experiments. Scale bars-10 µm. Statistical analysis was done using Mann-Whitney’s test (non-normalized graphs) (* p-value<0.05, ** p-value <0.01, *** p-value <0.001, **** p-value <0.0001, ns = not significant).

**Figure 6 Supplementary: (A)** Collapsed Z-stack images of stable adherent control WT-MEFs (CNT), WT-MEFs treated with siRNA against KIF5B (KIF5B KD), dynein (DYN KD) individually, or together (DUAL KD), immunostained with Giantin and Alpha tubulin (Alpha tub). Scale bars-10 µm.

