## Supplementary figures and images for "KIF5B and Dynein regulate adhesion-dependent Golgi organization and microtubule acetylation"

### Supp Figure 1

Figure 1 Supplementary

**(A)**

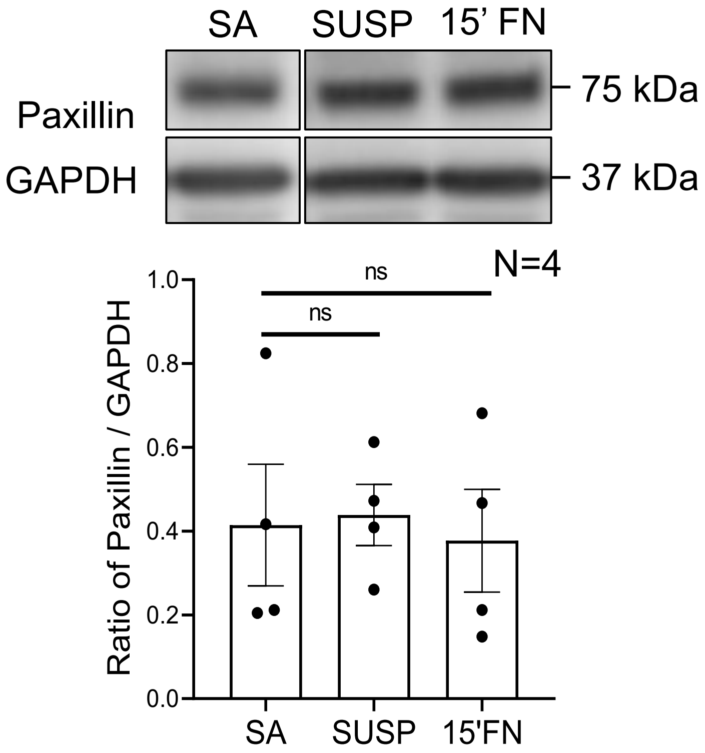

### Supp Figure 3

#### Figure 3 Supplementary

**(A)**

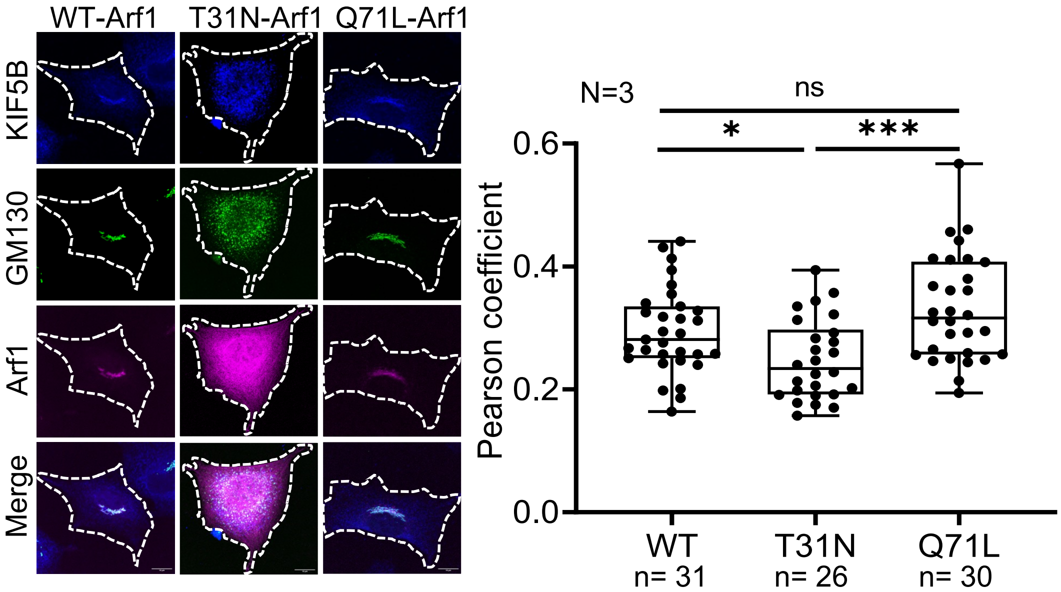

### Supp Figure 4

#### Figure 4 Supplementary

**(A)**

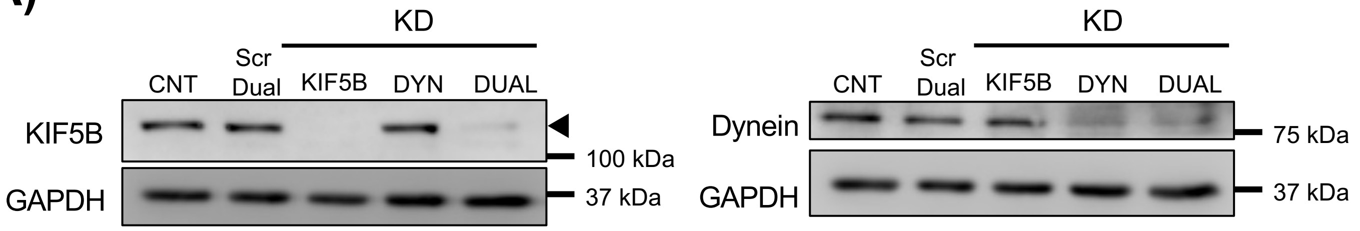


**(B)**

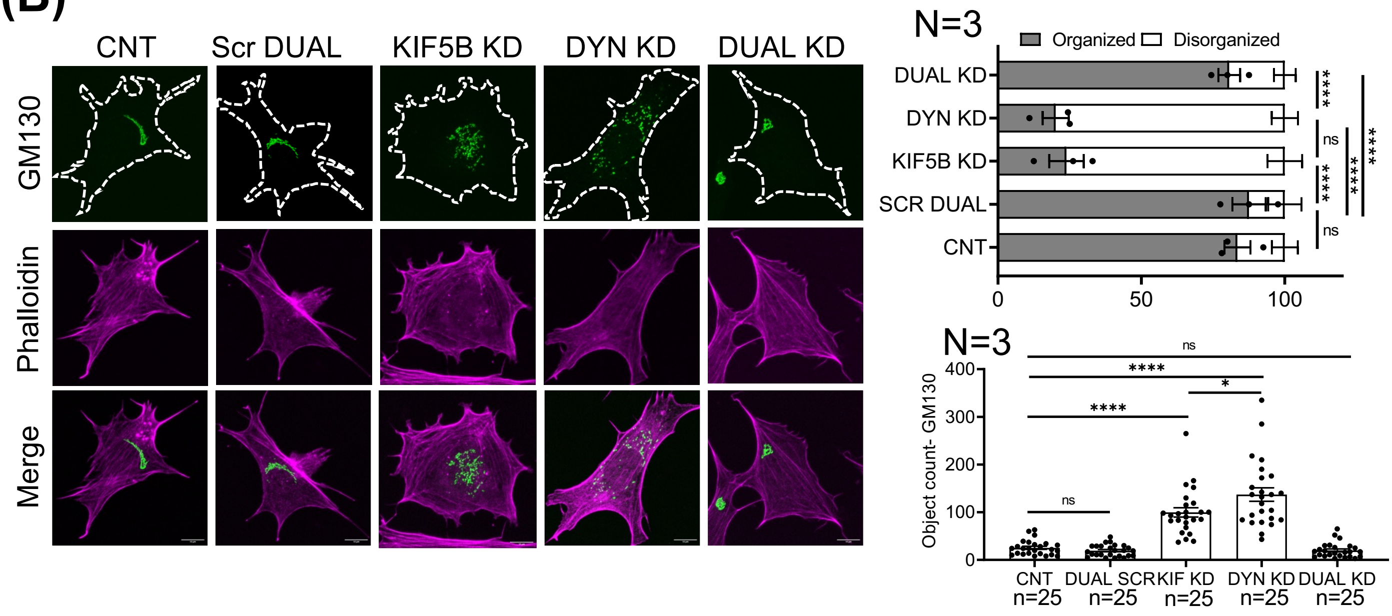


**(C)**

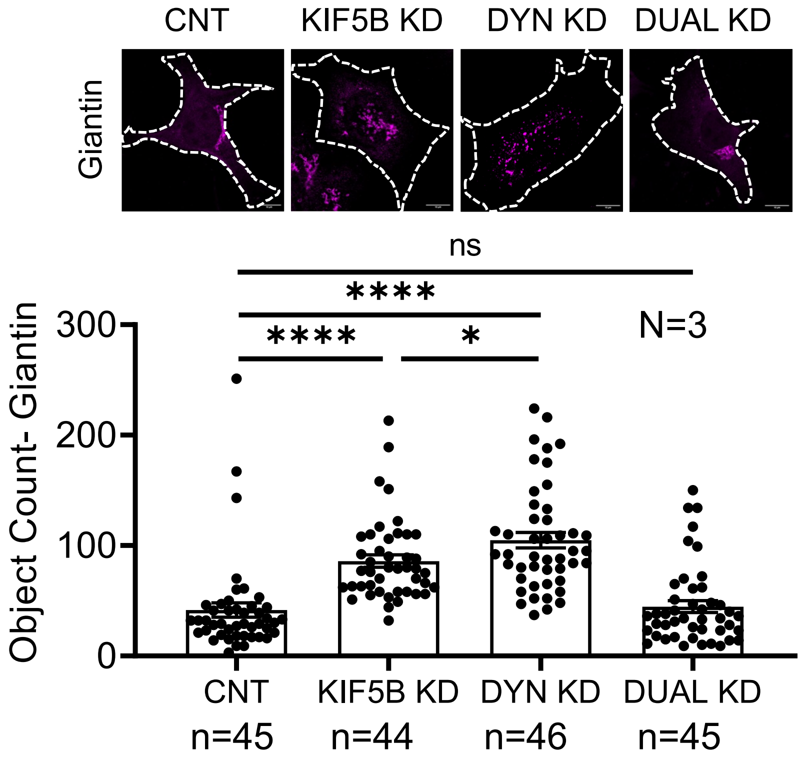

### Supp Figure 6

Figure 6 Supplementary

**(A)**

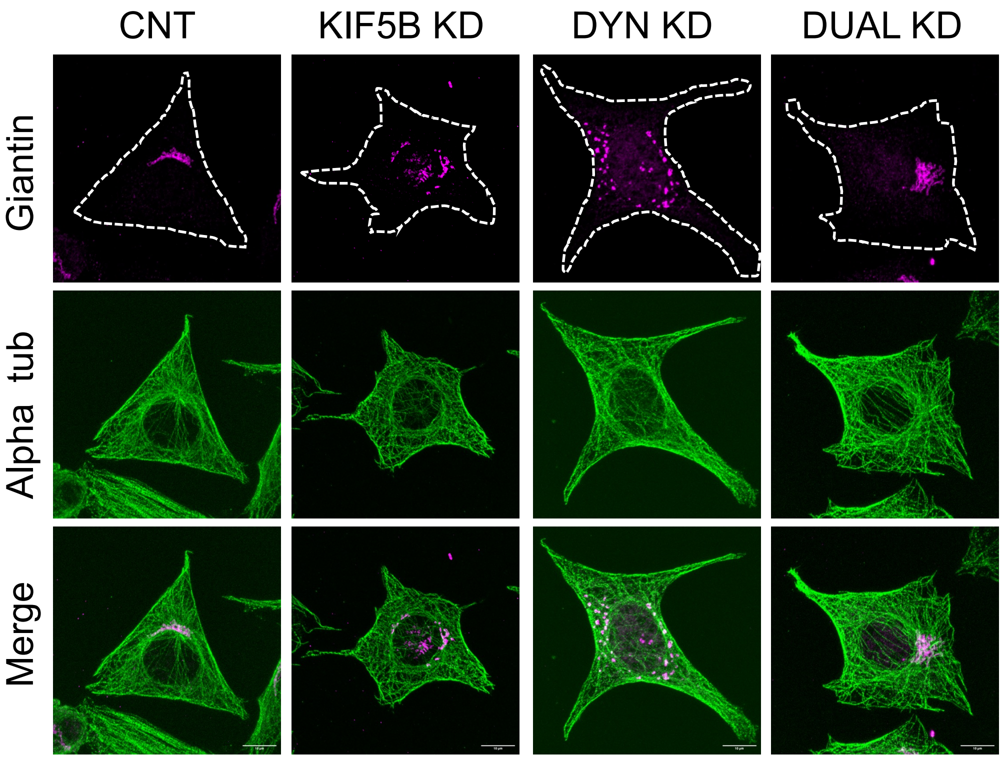
